# Glycosylation-modified antigens as a tolerance-inducing vaccine platform prevent anaphylaxis in a pre-clinical model of food allergy

**DOI:** 10.1101/2023.03.23.534004

**Authors:** Shijie Cao, Chitavi D. Maulloo, Michal M. Raczy, Matthew Sabados, Anna J. Slezak, Mindy Nguyen, Ani Solanki, Rachel P. Wallace, Ha-Na Shim, D. Scott Wilson, Jeffrey A. Hubbell

## Abstract

The only FDA-approved oral immunotherapy for a food allergy provides protection against accidental exposure to peanuts. However, this therapy often causes discomfort or side effects and requires long-term commitment. Better preventive and therapeutic solutions are urgently needed. We have developed a tolerance-inducing vaccine technology that utilizes glycosylation-modified antigens to induce antigen-specific non-responsiveness. The glycosylation-modified antigens were administered intravenously (i.v.) or subcutaneously (s.c.) and were found to traffic to the liver or lymph nodes, respectively, leading to preferential internalization by antigen-presenting cells, educating the immune system to respond in an innocuous way. In a mouse model of cow’s milk allergy, treatment with glycosylation-modified β- lactoglobulin (BLG) was effective in preventing the onset of allergy. In addition, s.c. administration of glycosylation-modified BLG showed superior safety and potential in treating existing allergies in combination with an anti-CD20 co-therapy. This platform may provide an antigen-specific immunomodulatory strategy to prevent and treat food allergies.

## Introduction

Food allergies are a growing concern, affecting more than 10% of the US population and showing a significant increase in prevalence in recent decades^1^. This active area of research has produced paradigm-shifting literature in the last decade alone, paving the way for novel FDA-approved treatments and NIAID-sponsored guidelines for preventative and therapeutic strategies. Palforzia, an FDA-approved oral immunotherapy (OIT) for peanut allergy, provides protection against accidental exposure to peanut. However, Palforzia and similar therapies often cause discomfort and side effects and are unlikely to induce long-lasting non-responsiveness in their current forms^2–4^. Studies such as LEAP (Learning Early About Peanut Allergy) and EAT (Enquiring About Tolerance) have produced promising and impactful insight into early introduction to prevent the onset of food allergies — exposing infants to major food allergens in their diet alongside breastfeeding leads to significantly reduced onset of food allergy compared to control subjects in five-year longitudinal studies^5, 6^. While early introduction guidelines are gradually being adopted by pediatricians and allergists across the US, one of the biggest barriers to effective prevention is compliance with the rigorous schedule of daily feeding of specific amounts of allergen to infants for years, as well as the risk of exposing other children in the household who might already be allergic^7^. Because of these limitations for both preventative and therapeutic approaches, alternatives that can provide lasting non-responsiveness or even tolerance to different allergens in a limited number of doses are needed.

Oral tolerance to food allergens is maintained through multiple mechanisms, with the gut-associated lymphoid tissues (GALT) playing a crucial role in controlling inflammatory immune responses to commensal bacteria and food proteins^8^. This process involves collection and processing of food antigens by antigen-presenting cells (APCs) and the induction of antigen-specific regulatory T cells (Tregs) or B cells (Bregs). Additionally, research has shown that the microbiome is linked to oral tolerance by stimulating the production of Tregs and suppressing type 2 T helper (Th2) immune responses^9–11^. A failure of oral tolerance can lead to the development of food allergy, caused by Th2-biased immune responses, which produce antibodies such as IgE and release inflammatory mediators, resulting in allergic reactions including anaphylaxis upon subsequent exposure to the allergens^12^.

OIT is an antigen-specific treatment that aims to desensitize patients to food allergens by orally administering small doses of the allergen in progressively larger amounts. However, OIT has limitations, such as common side effects including gastrointestinal symptoms, which can cause patients to discontinue treatment, and it may require long-term maintenance therapy^13, 14^. Alternatives such as subcutaneous or sublingual immunotherapies (SCIT or SLIT) have been studied, but they also have limitations, such as higher rates of anaphylactic reactions with SCIT or lower efficacy with SLIT^2, 15, 16^. To improve antigen-specific tolerance, researchers are investigating strategies such as modifying antigens through functional conjugation to polymers, nanoparticles, antibodies or small molecules that can improve the delivery of antigens or target specific immune cells^17^. Our lab has explored synthetic glycosylation of antigens to create a vaccine for inducing tolerance to antigens by leveraging delivery to C-type lectin receptors (CLRs) on APCs in the liver or in the peripheral lymph nodes (LNs)^18, 19^. Tolerance has been observed for three different glycosylations bearing either NAc-galactosamine (p(GalNAc)), NAc-glucosamine (p(GlcNAc)), or a new glycosylation strategy bearing mannose (p(Man), below. The glycosylation modification, when applied to gluten antigen in celiac disease, has safely progressed through a phase I clinical trial *(ClinicalTrials.gov Identifier: NCT04248855)*^20, 21^, showing that the platform was safe and reduced T cell responses following a gluten challenge; another phase I clinical trial in multiple sclerosis *(NCT04602390*) is underway. Our latest endeavors of applying synthetic glycosylation technology include preventing anti-drug antibody (ADA) responses to prolong the efficacy of immunogenic protein drugs^22^, and here we further applied this technology in preventing and treating food allergies.

The abundantly expressed mannose-binding receptors on diverse APCs function as excellent antigen internalization machinery and channel antigens to cross-presentational pathways^23, 24^. When paired with an adjuvant, a p(Man)-antigen conjugate is an effective vaccine^25, 26^, but we wanted to investigate the tolerogenic capacity of p(Man) when used in an unadjuvanted form. These glycosylation-modified antigens have resulted in the generation of robust antigen-specific CD4^+^ and CD8^+^ T cell tolerance and hypo-responsiveness to antigenic challenges via clonal deletion, anergy of activated T cells, and expansion of regulatory T cells^18, 19^. In this study, we sought to adapt the technology to modify food proteins and investigate the potential for preventing and treating food allergies through intravenous (for targeting liver), or subcutaneous (for targeting peripheral LNs) administration. Using a mouse model of cow’s milk allergy, we tested the efficacy and mechanisms of immune modulation of glycosylation-modified β-lactoglobulin (BLG), a major cow’s milk antigen, in both prophylactic and therapeutic settings.

## Results

### p(Man) delivered cow’s milk antigen to the liver or draining LNs through i.v. or s.c. administrations, respectively

Wallace et al. described the synthesis of a glycopolymer called p(Man), which effectively targeted the liver and reduced antigen-specific cellular and humoral responses to various immunogenic proteins^22^. Here, we conjugated the polymer to BLG using a self-immolative DBCO linker (**Figure 1A, Figure S1, S2**). Our conjugation method enables each antigen to be linked with multiple p(Man) polymers through the available primary amine groups. In the case of BLG, it contains 13 lysine residues, each providing a primary amine that could potentially undergo conjugation. However, we do not expect all these sites to participate in conjugation due to the constraints imposed by steric hindrance. As shown by SDS-polyacrylamide gel electrophoresis (SDS-PAGE), the conjugation of the DBCO linker to the free amine group of BLG resulted in an increase of molecular weight from 15 kDa to above 25 kDa. After the conjugation of p(Man) to the BLG-linker followed by purifying with size exclusion chromatography (SEC), the final product, BLG-p(Man), had an effective molecular weight of around 75-250 kDa (**Figure 1B, Figure S3**). The conjugated BLG-p(Man) also formed nanostructures with a measured diameter of 13.06 ± 0.45 nm (**Figure 1C**), The polydispersity index (PDI) was 0.46 ± 0.20, attributable to the variability in p(Man) conjugation to BLG, which lead to a spread in molecular weight (**Figure 1B**). The ζ-potential was measured at -2.63 ± 0.74, indicating a nearly neutral surface charge of the BLG-p(Man).

**Figure 1.**
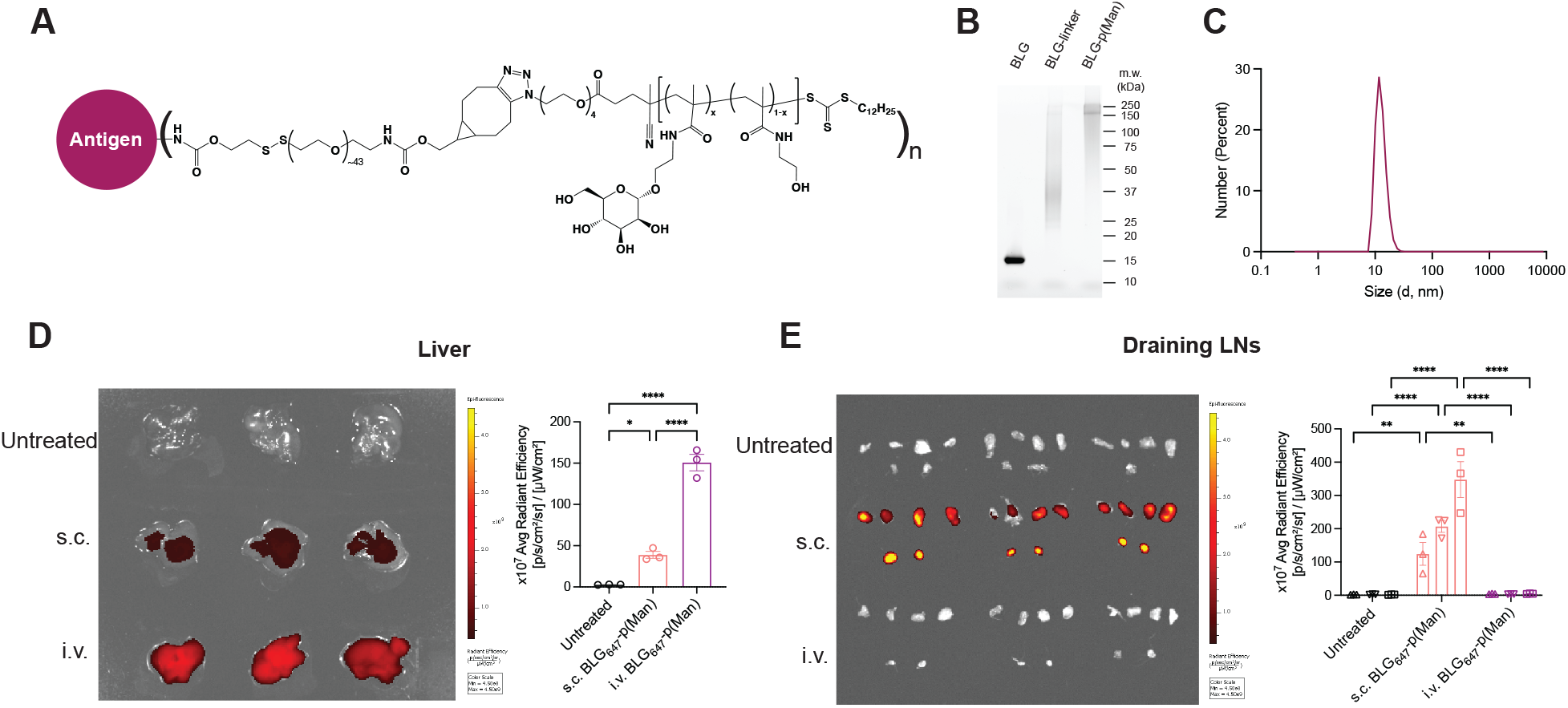
**A.** Chemical structure of antigen conjugated to p(Man) via a self-immolative linker. **B**. SDS-PAGE analysis of unmodified BLG, BLG-linker conjugate, and BLG-p(Man). **C**, Size distribution in terms of diameter of BLG-p(Man), measured by dynamic light scattering (DLS). **D, E.** IVIS imaging and analysis of fluorescence signals from mouse liver (**D**) and draining lymph nodes (top: two axillary and two brachial LNs; bottom: two popliteal LNs) (**E**). Mice were injected with BLG647-p(Man), through either i.v. or hock s.c. injections. Liver and draining LNs were harvested at 10 hr post-injection and imaged using IVIS. Fluorescence intensity was quantified as average radiant efficacy and compared by one-way ANOVA using Tukey’s *post hoc* test. Data represent mean ± s.e.m., *p < 0.05, **p<0.01, ***p<0.001, ****p<0.0001.

We have showed that glycosylation-modified ovalbumin (OVA, as a model antigen) is delivered to the liver through i.v. administration^18^ and to the injection site-draining peripheral LNs through s.c. administration^19^. To ascertain the biodistribution of the new glycosylation-modified antigen BLG-p(Man), we treated C57BL/6 mice with fluorescently labeled BLG647- p(Man) via tail vein i.v. or four hock s.c. administrations. After 10 hr, the mice were perfused, and the liver and hock-draining LNs (including axillary, brachial and popliteal LNs) were harvested for whole-organ fluorescence imaging (**Figure 1D, E**). The imaging showed that i.v. administration of BLG647-p(Man) preferentially targets the liver with a more than 3-fold increase compared to the same dose of BLG647-p(Man) injected subcutaneously. BLG647-p(Man) administered s.c. accumulated in the draining LNs, while a signal was not detectable in those LNs from the i.v. group.

### Intravenous administration of BLG-p(Man) prevents severe anaphylactic response to cow’s milk antigen in mice in a prophylactic model

Dysbiotic mouse models, including antibiotic-treated and gnotobiotic mice, have been used to gain insight and develop potential therapeutics for food allergies^27–29^. In this study, we used a dysbiotic mouse model utilizing C3H/HeJ mice, which are known for their susceptibility to food allergies due to their impaired TLR4 function^30, 31^. This model was utilized to investigate the potential of BLG-p(Man) in preventing food allergies (**Figure S4A**). We treated mice with an antibiotic cocktail one week prior to weaning and then added antibiotics to the drinking water throughout the study. At six days and three days before weaning, we injected mice i.v. with either PBS, 0.2 μg unmodified BLG per gram of body weight, or equivalent BLG-p(Man).

Starting at weaning, we sensitized mice using BLG plus cholera toxin (CT) as a mucosal adjuvant intragastrically (i.g.) once weekly for 5 weeks as described previously^7^. After five sensitizations, we challenged the mice intraperitoneally (i.p.) with BLG and evaluated anaphylaxis by measuring any drop in core body temperature for 90 min after the food challenge. We quantified serological markers, including BLG-specific IgE, total IgG, IgG1 and mouse mast cell protease 1 (mMCP-1). We observed that mice receiving either PBS or unmodified BLG were susceptible to anaphylactic responses after a BLG challenge, as evidenced by a drop in core body temperature (**Figure S4B**), and also produced BLG-specific IgG, IgG1 and IgE, as well as mMCP-1 (**Figure S4C-F**). In contrast, all mice treated with BLG-p(Man) were protected from an anaphylactic response to BLG sensitization and challenge, and there was minimal production of BLG-specific antibodies and mMCP-1. We further investigated cellular responses and cytokine production by restimulating mouse splenocytes with BLG *in vitro* (**Figure S4G, H**). We observed that cells from mice treated with BLG-p(Man) had a lower percentage of GATA3^+^ Th2 cells among CD4^+^ T cells upon 6 hr restimulation with BLG. The splenocytes from BLG-p(Man) pretreated mice also produced lower levels of cytokines upon restimulation, including IFNγ, IL-10 and IL-17A, compared to those from mice treated with PBS or unmodified BLG. For other cytokines including TNFα, IL-2, IL-4, and IL-13, we observed some upregulation from allergic mice that were treated with PBS or BLG, but not from mice treated with BLG-p(Man). These data suggest that prophylactic i.v. administration of BLG-p(Man) had promise in protecting mice from an anaphylactic response to BLG.

### Subcutaneous administration of p(Man) antigen conjugates provides an alternative route for inducing antigen specific tolerance

We have previously shown that glycosylation-modified conjugated antigens are successful at inducing both cellular and humoral tolerance through liver targeting, preventing the onset of autoimmune disease^18^ and ADA responses^22^. However, s.c. administration could be clinically preferable for preventing food allergy in infants and children and may present fewer systemic safety concerns if administered to patients with existing food allergies. Our lab has recently demonstrated that p(GlcNAc)-conjugated antigen can lead to similar tolerogenic results when targeted to LNs through s.c. administration^19^. APCs in the LNs share several CLRs and scavenger receptors with hepatic APCs^32^, suggesting that the superior uptake efficiency and subsequent accumulation of glycosylation-modified antigen seen in the liver could be replicated in the LN microenvironment.

To investigate tolerance induction by treatment with p(Man)-conjugated antigens when administered s.c., we used OVA as a model protein in an OVA-reactive transgenic T cell receptor (TCR) mouse model (**Figure 2A**). The p(Man) was conjugated to OVA (OVA-p(Man)) through the self-immolative linker as described previously^18, 19^. On day 0, we adoptively transferred CD45.1^+^ OVA-specific CD8^+^ and CD4^+^ T cells (OTI and OTII cells, respectively) into recipient CD45.2^+^ C57BL/6 mice. On days 1 and 7, we injected mice s.c. in four hocks with OVA-p(Man), unmodified OVA, or saline. We then immunized mice with OVA and LPS on day 16 and assessed the OVA-specific immune responses after sacrificing mice on day 21. We examined the phenotype changes in either OTII or OTI cells to demonstrate antigen-specific tolerance. These cells respond specifically to the OVA, which we injected subcutaneously into the mice. From the hock-draining LNs (axillary, brachial and popliteal LNs), we observed a lower percentage of OVA-specific OTI (15%) and OTII cells (2%) of total CD4^+^ T cells from mice treated with OVA-p(Man), compared to mice treated with saline (35% for OTI and 14% for OTII) (**Figure 2B-D**). From the OTII cells, the OVA-p(Man) treatment induced more antigen-specific Foxp3^+^CD25^+^ regulatory T cells (Tregs) with a 2-fold increase compared to the free OVA-treated mice (**Figure 3A, Figure S5**). From the OTI cells, we observed significantly higher levels of co-inhibitory markers, including PD-1 and LAG-3, from the OVA-p(Man) group (**Figure 3B**). Upon restimulation of lymphocytes from hock-draining LNs with their cognate peptide OVA257-264 *ex vivo*, a smaller fraction of OTI cells from mice treated with OVA-p(Man) produced pro-inflammatory cytokines including IFNγ and TNFα (**Figure 3C, Figure S6A**). Similarly, a significantly lower fraction of OTII cells produced IFNγ and TNFα upon restimulation with their cognate OVA323-339 peptide (**Figure 3D, Figure S6B**). We also incubated the cells isolated from hock-draining LNs with whole unmodified OVA protein for 3 days and measured cytokine secretion. The cells from mice treated with OVA-p(Man) showed a significant reduction in pro-inflammatory cytokines including IFNγ, TNFα, IL-2, IL-17A, and IL-22 upon restimulation, compared to cells from saline-treated mice (**Figure 3E**). Thus, we confirmed that s.c. delivery of antigens via p(Man) results in antigen-specific tolerance, characterized by antigen-specific T cell deletion, upregulation of co-inhibitory markers, induction of Tregs, and an abrogation of pro-inflammatory cytokines upon antigenic challenge and restimulation.

**Figure 2.**
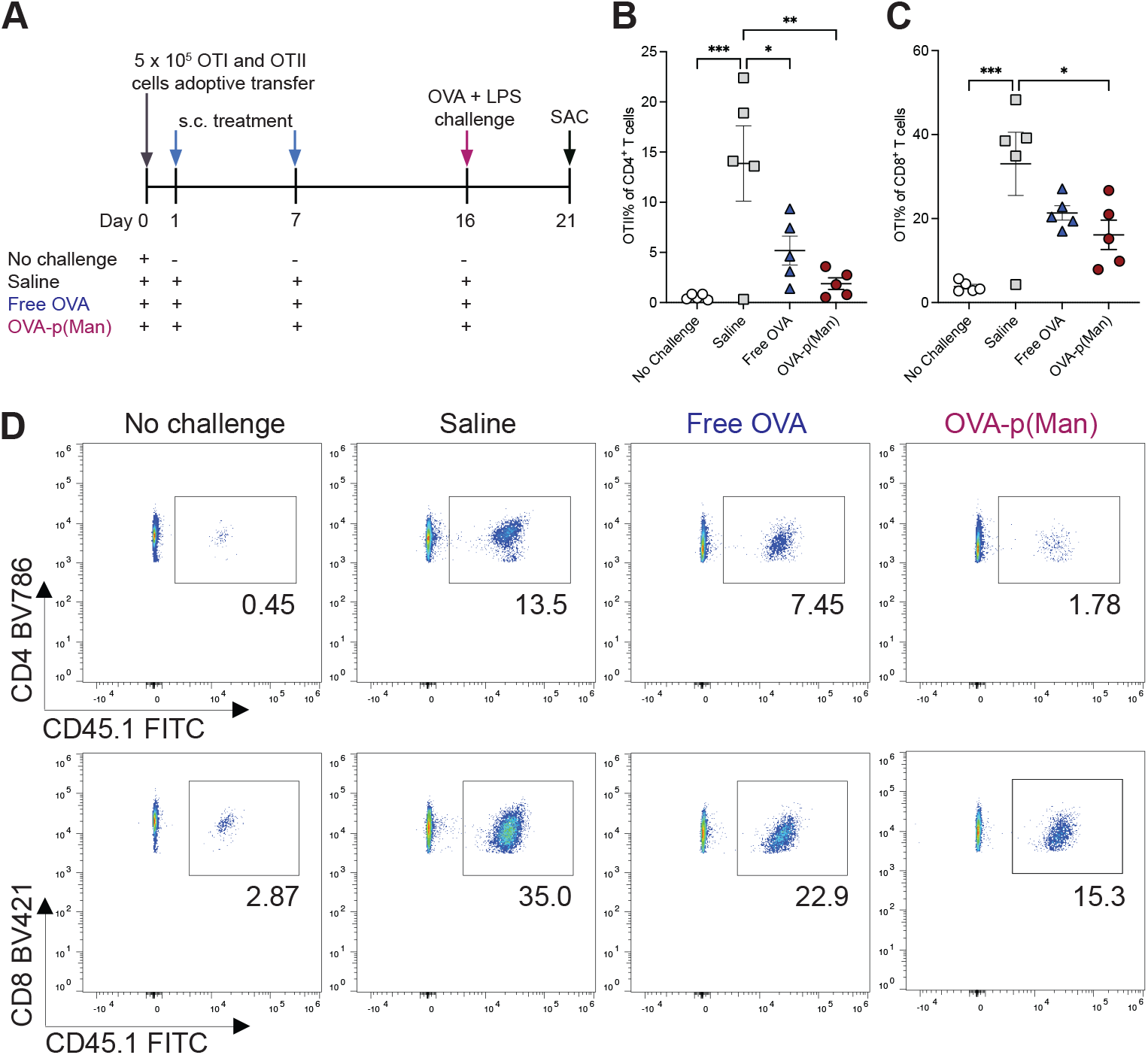
p(Man)-conjugated antigens administered s.c. induce OTI and OTII cell deletion to antigen challenge. **A.** Experimental timeline of prophylactic tolerance. C57BL/6 mice that had received an adoptive transfer of OTI and OTII cells were treated on days 1 and 7 with saline or 20 μg of OVA as unmodified OVA, or OVA-p(Man) s.c. in four hocks. On day 16, the mice in all treatment groups were challenged with OVA + LPS in the hocks, then 5 days later the injection site-draining LNs (dLNs) were examined for an OVA-specific immune response. **B, C.** Quantification of the percentage of OTII (C) and OTI (D) cells of total CD4^+^ or CD8^+^ cells, respectively, in the dLNs on day 21. **D.** Representative flow cytometry plots of percentage of OTII (top) and OTI (bottom) cells. Data represent mean ± s.e.m.. Statistical differences were determined by one-way ANOVA using Dunnett’s *post hoc* test (*p < 0.05, **p<0.01, ***p<0.001, ****p<0.0001).

**Figure 3.**
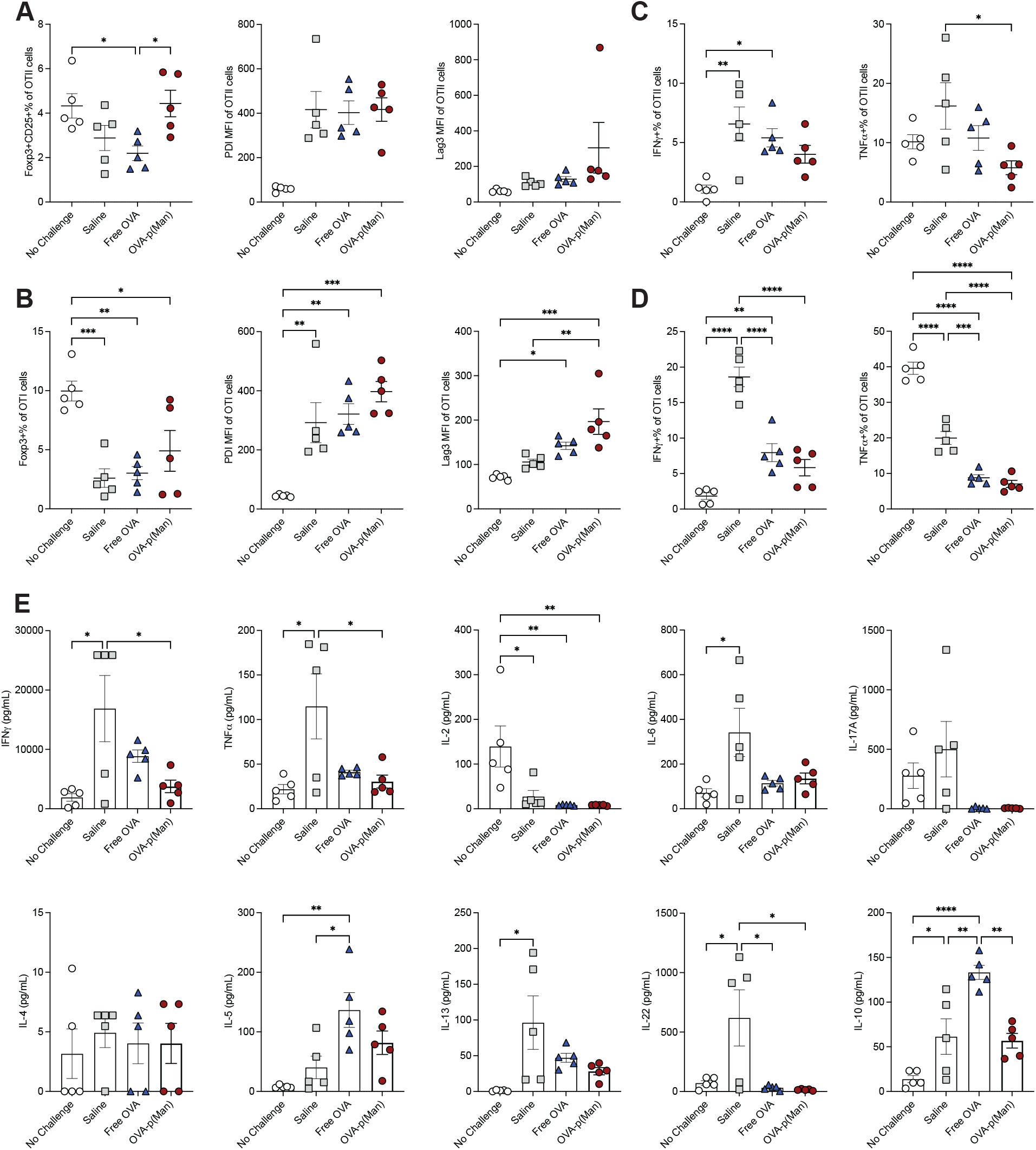
p(Man)-conjugated antigens administered s.c. CD8^+^ and CD4^+^ antigen-specific T cell tolerance to antigen challenge, from experiment in Fig. 2. **A, B.** The percentage of Foxp3^+^CD25^+^ cells, and the mean fluorescent intensity (MFI) of PD1 and Lag3 expression of OTII (A) or OTI (B) cells from mice dLNs at day 21. **C**. Percentage of IFNγ^+^ and TNF𝛂^+^ cells of OTII cells upon restimulation with OVA323-339 peptide. **D**. Percentage of IFNγ^+^ and TNF𝛂^+^ cells of OTI cells upon restimulation with OVA257-267 peptide. **E.** Production of cytokines by isolated cells from dLNs upon restimulation with OVA protein for 3 days. Data represent mean ± s.e.m.. Statistical differences were determined by one-way ANOVA using Dunnett’s *post hoc* test (*p < 0.05, **p<0.01, ***p<0.001, ****p<0.0001).

### Subcutaneous administration of glycosylation-modified BLG prevents the onset of BLG allergy through inhibiting the generation of BLG-specific antibodies and Th2 responses

Based upon the results with model protein OVA, we sought to replace i.v. administration of BLG-p(Man) with s.c. administration. The previous animal model of BLG allergy (**Figure S4**) presented limitations, as the mice were not consistently induced with BLG allergy despite receiving the same dose of BLG and CT at the same times. In this experiment, we modified the protocol by optimizing the timing and the doses of BLG and CT, resulting in more consistent and robust allergic responses after sensitization.

In this new model, the mice were kept on an antibiotic cocktail drinking water regimen starting after weaning for the duration of the experiment. One day after weaning, mice were s.c. administered twice, 6 days apart, with either saline or an equivalent amount of 1 µg/g body weight BLG in the form of free BLG or BLG-p(Man). Three days after the second administration, all mice were intragastrically sensitized weekly for 5 weeks with BLG plus CT. One week after sensitization, all the mice were challenged by oral gavage with BLG, and their core body temperature was recorded to evaluate their allergic responses. Compared to the saline-treated mice, mice pre-treated with BLG-p(Man) experienced significantly reduced anaphylactic drops in core body temperature (**Figure 4B**). While mice treated with free BLG did not experience as drastic a drop in body temperature compared to mice treated with saline, there was no significant difference between saline-treated mice and free BLG-treated mice. We collected blood after sensitization to assess the generation of BLG-specific antibodies in the serum.

**Figure 4.**
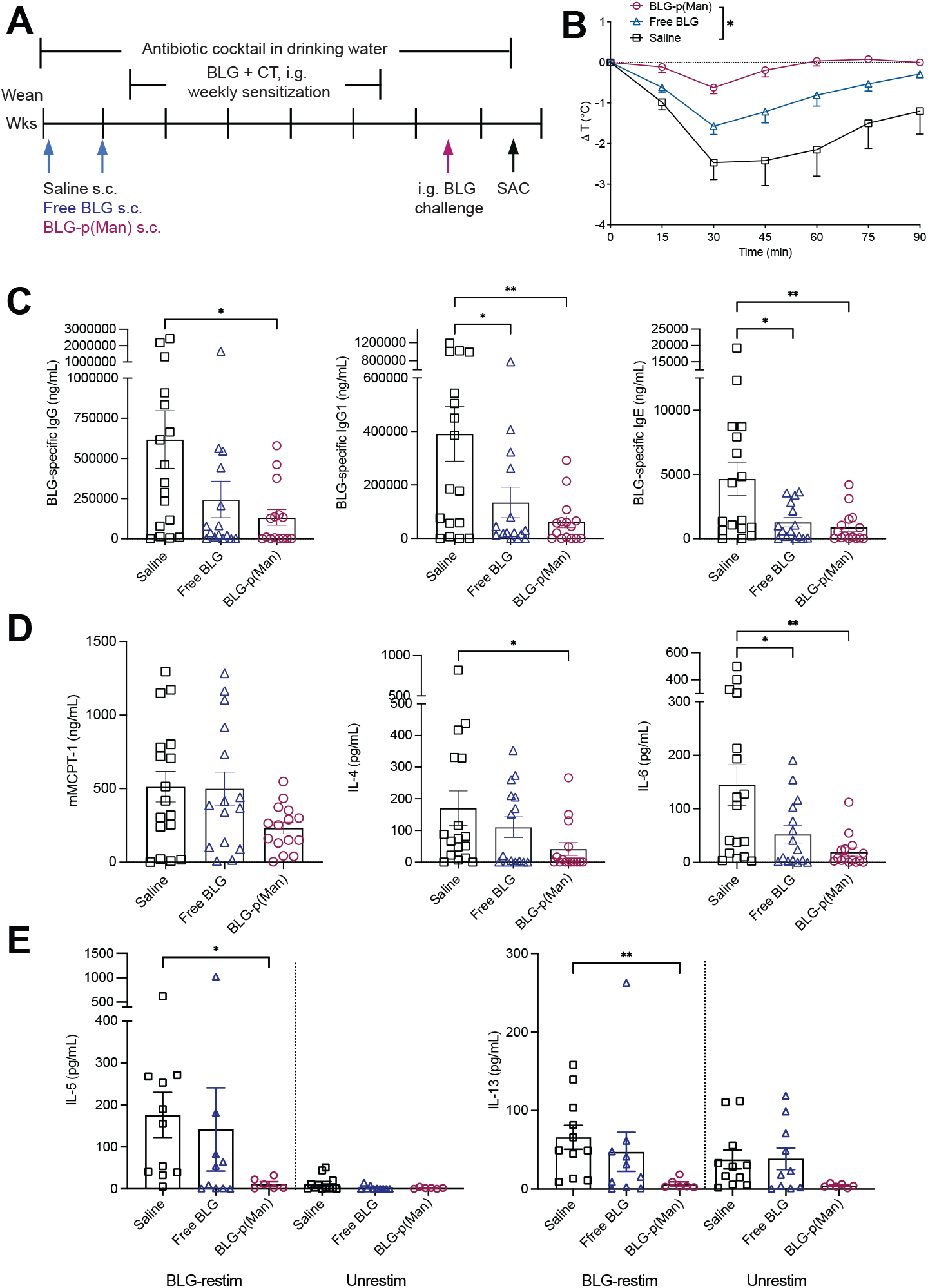
p(Man)-conjugated BLG protects mice from exhibiting allergic response to cow’s milk antigen. **a.** Experimental design. C3H/HeJ mice were given antibiotic water throughout the experiments and were s.c. administered with two doses of either saline, unmodified BLG, or BLG-p(Man) before sensitization. The mice were then orally challenged with BLG to assess the allergic reactions. **B,** Change in core body temperature at indicated time points following challenge with BLG in BLG + CT sensitized mice. **C.** BLG-specific IgG, IgG1, and IgE in mice serum collected three days before the oral challenge. **D.** mMCPT-1, IL-6, and IL-4 from post-challenge serum of mice in B. **E.** Th2 cytokine (IL-5 and IL-13) levels from culture supernatants of splenocytes from mice sacrificed one week post-challenge and stimulated for 4 days with BLG. Data were pooled from three experiments. Data represent mean ± s.e.m.. Statistical differences determined by one-way ANOVA using Dunnett’s post hoc test (B-D) or Kruskal-Wallis test (E). (*p < 0.05, **p<0.01).

Compared to saline-treated mice, we observed that all BLG formulations, including the free BLG and BLG-p(Man), significantly inhibited the generation of BLG-specific IgG1 and IgE (**Figure 4C**). The BLG-p(Man) treated mice also showed a significantly lower level of BLG-specific IgG. From the blood collected at 1.5 hr post-challenge, we observed that mice treated with BLG-p(Man) had reduced levels of mMCPT-1 (**Figure 4D**), which is released by mucosal mast cells upon degranulation during allergic hypersensitivity responses^33, 34^. IL-6 and IL-4 have been found to be elevated during anaphylaxis in food allergic patients^35^. The mice treated with saline or free BLG also showed elevated IL-6 and IL-4 levels in the serum upon BLG challenge, but not the mice treated with BLG-p(Man) (**Figure 4D**). We further investigated cytokine production by restimulating splenocytes with BLG *ex vivo* (**Figure 4E**). We observed that cells from sensitized mice treated with saline or free BLG had upregulated Th2 cytokines including IL-4 and IL-13, but not from mice treated with BLG-p(Man).

In addition to p(Man), we also tested the two other synthetic glycopolymers, p(GalNAc) and p(GlcNAc), developed in our lab, by conjugating them to BLG and using them in the same prophylactic model of BLG allergy (**Figure S7**). We observed similar protection from an anaphylactic response to BLG and suppression of antigen-specific Th2 responses and humoral antigen-specific antibody responses with BLG-p(GalNAc) and BLG-p(GlcNAc) treatments as we observed with BLG-p(Man). This suggests that all these glycosylation-modified BLG conjugates can effectively protect against an anaphylactic response to BLG.

### BLG-p(Man) demonstrates superior safety compared to free BLG when injected into allergic mice and shows potential in reversing existing allergies

SCIT is FDA-approved for treating reactions to various allergens such as trees, grass, house dust and insect stings, but not yet for food allergens^36, 37^. Although SCIT has shown promising efficacy in a peanut allergy clinical trial, systemic side effects (e.g. pulmonary and gastrointestinal symptoms) were reported that deterred further SCIT testing and favored the development of epicutaneous strategies instead^38, 39^. With the evidence that glycosylation-modified BLG successfully protected mice from developing cow’s milk allergy, we aimed to investigate if administration of glycosylation-modified food allergens to sensitized mice could provide a safe and effective way to treat existing food allergies. In our initial attempts, we injected three escalating doses of either saline, BLG-p(Man), or free BLG subcutaneously into four hocks of mice that had been sensitized to BLG (**Figure S8A**). We observed that the mice treated with free BLG began to show anaphylactic reactions right after the first injection and experienced more severe reactions after the second injection, with more than half of the mice dying from free BLG treatment (**Figure S8B**). In contrast, mice that received BLG-p(Man) at the equivalent BLG dose remained healthy after the injections. We further investigated cytokine production by restimulating cells isolated from spleen and mesenteric LNs, and we observed that mice treated with BLG-p(Man) showed significantly reduced Th2 cytokine production in both organs (**Figure S8F, G**). This suggests that Th2 suppression by the glycosylation-modified antigens was effective not only in the prophylactic setting but also in therapeutic settings in already-allergic mice. However, in this experiment we did not observe a significant difference in the core body temperature drop upon BLG challenge between the two groups (**Figure S8C**).

This may be attributed to the presence of existing humoral responses in the sensitized mice, such as IgE-expressing memory B cells, plasmablasts, and long-lived IgE-producing plasma cells^40–43^, which may not be targeted by the BLG-p(Man) treatment.

Given the lack of strategies that can specifically deplete the plasma cells with minimal side effects *in vivo*, we incorporated a general B-cell depletion method using anti-CD20 monoclonal antibodies (mAbs). Anti-CD20 mAbs such as rituximab and obinutuzumab are FDA-approved and used in clinic for treating diseases such as non-Hodgkin’s lymphoma (NHL) and rheumatoid arthritis (RA)^44^. Here, we injected mice with a single dose of anti-CD20 mAbs (clone mb20-11), followed by two s.c. injections of BLG-p(Man) (**Figure 5A**). We found that a single i.p. injection of anti-CD20 mAb efficiently depleted B220^+^CD19^+^ B cells and B220^+^CD138^hi^ plasmablasts for up to 30 days, which covers the duration of the treatments and the first oral BLG challenge (**Figure S9, Figure S10**). The BLG-p(Man) treatment was safe to the mice, as there was no significant drop in core body temperature within 30 min after receiving the treatment (**Figure 5B**), while the second injection of free BLG still caused severe anaphylactic reactions and lethargy in more than half of mice in that group. Two weeks after the therapy, we challenged the mice and observed ameliorated allergic responses with the BLG-p(Man) treatment compared to the control group, measured by core body temperature during the oral BLG challenge (**Figure 5C**) and mMCPT-1 levels in their post-challenge serum (**Figure 5D**).

**Figure 5.**
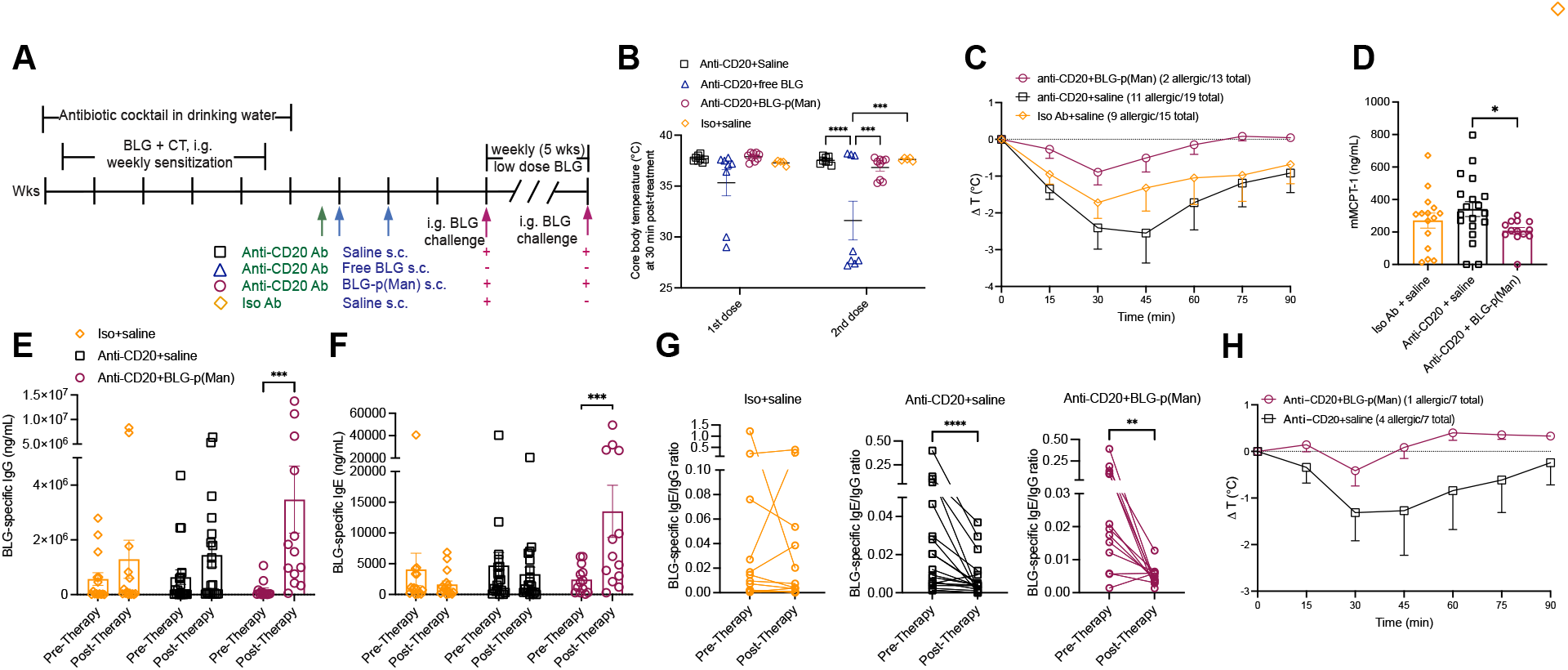
BLG-p(Man) treatment was safe for allergic mice and protected against severe anaphylaxis upon BLG challenge. **A.** Experimental design. C3H/HeJ mice were given antibiotic water and sensitized weekly by intragastric gavage of 0.1 mg/g body weight of BLG plus 6.7 mg/g body weight of the mucosal adjuvant cholera toxin. 5 days after sensitization, the mice were i.p. injected with anti-CD20 antibody or isotype control, and two days later followed by s.c. administration of two doses of saline, free BLG, or BLG-p(Man). The mice were then challenged i.g. with BLG to assess their anaphylactic responses (except for the free BLG group, in which more than half of mice died from anaphylactic reactions upon treatment). Mice treated with anti-CD20 and saline, or BLG-p(Man) were given weekly low doses of BLG (1 mg per mouse) and challenged again afterwards. Allergic mice were defined as core body temperature dropping by 1°C or more upon BLG challenge. **B**. The core body temperature at 30 min after each therapeutic injection. **C.** Change in core body temperature at indicated time points following oral challenge with BLG. **D.** mMCPT-1 measured from post-challenge serum of mice in C**. E, F.** BLG-specific IgG (E) and IgE (F) in the mouse serum before and after treatment. **G.** Change of the ratio of BLG-specific IgG to IgE before and after the treatment. Data were pooled from three experiments in C-G. **H.** Change in core body temperature of mice kept in weekly dosing of BLG after treatment, fol212lowing a second oral challenge with BLG. Data represent mean ± s.e.m.. Statistical differences determined by one-way ANOVA using Dunnett’s post hoc test (B), Kruskal-Wallis test (D), two-way ANOVA using Bonferroni’s post hoc test (E, F), or Wilcoxon test (G) (*p < 0.05, **p<0.01, ***p<0.001, ****p<0.0001).

Although not all mice responded to the oral BLG challenge in these experiments, the anti-CD20 mAb plus BLG-p(Man) treatment was able to reduce the number of allergic mice to 2 out of 13, while over half of the mice in the other two treatment groups were still allergic. Contrary to our results in the prophylactic setting, we observed that the BLG-p(Man) treatment significantly increased BLG-specific IgG and IgE levels in sensitized mice (**Figure 5E, F**). While IgE antibodies bind to mast cells via FcεRI and trigger hypersensitivity upon antigen encounter, antigen-specific IgG has been shown to promote food allergen tolerance and suppress existing Th2 and IgE responses through FcγRIIb signaling^45, 46^. In this experiment, we observed that BLG-specific IgG was increased much more (181-fold increase) than the BLG-specific IgE (7-fold increase), which might provide a mechanism of protection against anaphylactic reactions in the therapeutic setting. Furthermore, we compared the ratio of BLG-specific IgE to IgG, higher ratios being an indicator of a bias towards allergic responses instead of protection (**Figure 5G**)^47^. The isotype mAb treatment did not change the ratio of IgE to IgG, however, the anti- CD20 mAb treatment resulted in significant reduction of the IgE/IgG ratio. Moreover, the subsequent BLG-p(Man) treatment reduced the ratio from an average of 0.078 to 0.0050 (15.6- fold reduction), while the subsequent saline treatment (i.e., anti-CD20 mAb alone) reduced the ratio from 0.047 to 0.0076 (6.2-fold reduction).

Antigen-specific IgGs have been shown to increase, regardless of their subtypes, in response to OIT in both preclinical and clinical studies^46–48^. This can be beneficial in neutralizing allergens, as well as blocking IgE-mediated responses by binding to mast cells and basophils^49^. In C3H mice, IgG2a — a Th1-biased antibody subtype — stands out for its higher increase to various immunotherapies^50, 52, 54, 56^, potentially through the suppression of Th2-driven allergic responses. We further investigate the changes in BLG-specific IgG2a and IgG1 subtypes in response to BLG-p(Man) therapy in allergic mice. We observed that IgG2a levels were significantly increased in mice serum after BLG-p(Man) treatment (**Figure S11A**), while the increase in IgG1 — a Th2-biased antibody subtype — was not significant (**Figure S11B**). Mice treated with only anti-CD20 Ab or with saline did not experience any changes in these IgG subtypes.

After the first oral challenge of BLG, we kept two groups of mice that had been treated with either saline or BLG-p(Man) on weekly dietary doses of BLG for 5 weeks, and we gave them a second oral challenge to evaluate the durability of treatment, especially when the B cells were reconstituted one month post-anti-CD20 mAb treatment. After the dietary doses, there was only 1 out of 7 mice treated with anti-CD20 mAb followed by BLG-p(Man) that showed mild allergic reactions to the oral BLG challenge, while 4 out of 7 mice in the group treated with anti- CD20 mAb followed by saline showed anaphylactic responses (**Figure 5H**). This result demonstrates the potential of glycosylation-modified BLG in treating existing cow’s milk allergy when used in combination with B cell depletion.

## Discussion

Antigen-specific therapies are on the rise as we gain a better understanding of the underlying molecular mechanisms of different approaches. OIT for food allergies, as an antigen-specific treatment, requires long-term commitment to achieve the desensitization goal and is associated with safety issues including gastrointestinal hypersensitivity and risk of anaphylaxis even after the maintenance dose has been reached^13^. In this study, we investigated the use of glycosylation-modified antigens as a tolerance-inducing vaccine platform to prevent and treat cow’s milk allergy in mice. The glycosylation-modified antigens target key immunomodulatory sites in the body, such as the liver and lymph nodes, via i.v. or s.c. administration routes, respectively, and are preferentially internalized by carbohydrate-binding receptors on APCs^18, 19^. The antigen is released in the endosomal environment via a self-immolative linker for further processing and presentation by APCs, which then educates the immune system to respond to these antigens in an innocuous way.

In a prophylactic setting, we demonstrated that pretreating mice with glycosylation-modified BLG prior to sensitization prevented allergic responses to an oral BLG challenge: the pretreated mice were resistant to drops in their core body temperature characteristic of anaphylaxis, and experienced lower levels of markers of allergic reaction and inflammation including mMCPT-1, IL-4, and IL-6 in their post-challenge serum. Additionally, the treatment resulted in modulation of both humoral and cellular compartments, demonstrated by the inhibition of BLG-specific IgG, IgG1, and IgE antibodies, and BLG-specific Th2 responses. Our group has previously investigated mechanisms of prophylactic antigen tolerance induced by glycosylation-modified antigens administered via both i.v. and s.c. routes^18, 19, 22^. I.v. administered glycosylation-modified antigen targets hepatic APCs resulting in antigen-specific CD4^+^ and CD8^+^ T cell depletion and anergy, as well as Treg induction; s.c. administered glycoantigen targets similar scavenger receptors but on lymph node-resident dendritic cells to achieve antigen hyporesponsiveness via similar axes of T cell tolerance in addition to LAG-3 signaling. Results of s.c. OVA-p(Man) prophylactic tolerance conducted in the model antigen OVA transgenic model in this study revealed similar mechanisms at play, as shown by antigen-specific deletion, upregulation of LAG-3, Treg induction and deep reduction of inflammatory cytokines upon antigen challenge. Importantly, Th2 cytokines (IL-4, IL-13) were impacted, providing early evidence for humoral tolerance, and a reduction in T-cell may help suppress the downstream cascade of B cell activation and antibody production, consistent with our observations in the BLG challenge studies.

While prophylactic efficacy in the form of a tolerance-inducing vaccine serves as a good proof of concept and mechanistic validation, a more clinically useful strategy would be reducing allergic responses in already-allergic individuals. Indeed, studies revealed that around 7.6% of children and 10.8% of adults in the United States have probable food allergies, and the number is still on the rise^1, 51, 53^. Here, a crucial consideration for potential therapeutics is safety, as exposure to even a tiny amount of allergy-causing food in a sensitized individual can rapidly propagate a potentially deadly anaphylactic response. Our lab has generated evidence that the glycopolymer may act as a steric shield, protecting the antigen from antibody recognition and opsonization upon systemic administration, and it is reasonable to extrapolate these findings to the food allergy context where the glycopolymer shield could prevent recognition by sensitized mast cells and basophils, abrogating the first step in an allergic reaction. Nevertheless, due to the inherent risk factors that could deter adoption by clinicians and allergic patients, we chose to evaluate the therapeutic efficacy of BLG-p(Man) only when s.c. administered, thus providing a safer and more convenient local route of entry and a more convenient practice. As anticipated, two s.c. injections of BLG-p(Man) were safe and did not result in a temperature drop in mice that received them, whereas mice that received two doses of unmodified BLG experienced a significant temperature drop (**Figure 5B**). SCIT for food allergies has previously been discontinued due to safety concerns^38, 39^. Current SCIT treatments, used for conditions including allergic rhinitis and asthma, employ unmodified antigen and are associated with adverse events including pulmonary and gastrointestinal symptoms^36, 55^. Our results suggest that modifying SCIT antigens, in this case through glycosylation, can significantly enhance their safety and tolerability profile compared to unmodified antigens. As shown in **Figure S8B**, a third s.c. injection of BLG-p(Man) (10 µg) did result in some reactogenicity and a temperature drop. Thus, while this study provides a good starting point towards identifying an optimal dosing and therapeutic window, the dosing strategy needs to be thoroughly optimized to better balance the need for safety against improved efficacy to maximize therapeutic benefit.

To overcome the limitations of pre-existing humoral immunity, we introduced a co-therapy with anti-CD20 mAb to deplete B cells. Pre-treatment with anti-CD20 depleted ∼99% CD19^+^B220^+^ B cells in under a week and was sustained for 30 days (**Figure S9**). Subsequent glycosylation-modified BLG therapy resulted in ameliorated anaphylactic responses, along with an increase in BLG-specific antibody levels and a decrease in the ratio of IgE to IgG: We observed that anti-CD20 mAb treatment significantly reduced the BLG-specific IgE/IgG ratio, and subsequent SCIT treatment with BLG-p(Man) further lowered it by 34% (**Figure 5G**). This is consistent with the profile of clinically approved OIT treatments and provides a mechanistic basis for the ameliorated anaphylactic responses after oral BLG challenge. Studies have shown that OIT induces strong IgG responses, which encompasses all IgG subclasses, but only modest reductions in specific IgE antibodies^46, 47, 57–59^. Moreover, these allergen-specific IgGs effectively neutralize allergens and also compete for IgE binding *via* interactions with FcγRIIB, thus suppressing IgE-mediated effector cell activation^60, 61^. The BLG-p(Man) treatment substantially increased the BLG-specific IgG levels, which outcompeted the increase in IgE, resulting in a reduced IgE/IgG ratio (**Figure 5E-G**). Interesting, we also observed that the BLG-p(Man) treatment significantly induced BLG-specific IgG2a production, an Th1-biased IgG subtype that can be particularly protective against Th2-biased allergic responses in mice^50, 52, 54, 56^. This suggests a specific Th2 suppression mechanism induced by BLG-p(Man) therapy, which was further supported by several *in vitro* antigen restimulation cytokine analyses (Figure 4D, E; Figure S4G, S7D, E, and S8 F,H).

Durability of treatment is an important differentiator in the food allergy therapeutic landscape. The only FDA-approved OIT, Palforzia, requires 14 dose escalation treatments, including several administrations at each dose level before a maintenance dose can be attained, which translates to a cumbersome dosing paradigm and a potential lifelong treatment for efficacy. Here, we show that two therapeutic doses of BLG-p(Man) were sufficient for generating durable therapeutic benefit, as only 1/7 treated mice experienced an allergic response after receiving the long-term challenge, compared to 4/7 saline-treated mice (**Figure 5H**). Treatment durability was achieved 5 weeks following the first oral challenge and extended ∼4 weeks beyond B cell reconstitution (**Figure S9**), suggesting that repeated anti-CD20 administration might not be required, minimizing potential immunosuppressive adverse events such as severe infections resulting from its use^62^. We have previously observed that the tolerance signatures induced by treatment with glycosylation-modified antigens occur at the transcriptional level and result in a durable effect^22^. Even when B cell populations return post-anti-CD20 treatment, this durable effect persists, which may lessen the need for multiple doses of anti-CD20 in our combination therapy.

In a therapeutic context, this study shows the combinational effect of B cell depletion coupled with immunological reprogramming, here though administration of a glycosylation-modified antigen. Anti-CD20 or other B cell depletion therapies are currently not approved in the context of food allergies but are being investigated in the context of inflammatory and autoimmune indications, with some degree of success^44, 63^. We envision that our findings would spur further investigations to refine this strategy, identify suitable patient populations, and determine the optimal circumstances under which a similar approach could be clinically viable.

Alternative steps consist of further investigating combination strategies with other depleting agents capable of more directed or targeted depletion and that present a lower barrier to entry from a translational perspective. For example, an ongoing study called Omalizumab as Monotherapy and as Adjunct Therapy to Multi-Allergen Oral Immunotherapy in Food Allergy Participants (OUtMATCH, *NCT03881696*) aims to investigate the efficacy of anti-IgE therapy (omalizumab) in participants who are allergic to peanut and at least two other foods.

Our work highlights the synthesis of several glycopolymers that can be conjugated to free amines on proteins using a self-immolative, endosomally cleavable linker. The versatility and mild conditions of the conjugation chemistry ensure that the strategy can be universally applied to any protein food allergen that contains a native or engineered primary amine.

Additionally, the customizable platform allows for the treatment of patients against multiple allergies with infrequent dosing. For preventive purposes, this tolerance-inducing vaccine might be appropriate in infants who display atopic symptoms, such as eczema, that put them at higher risk and are currently used clinically as risk factors to identify infants who should receive early introduction^64^. In this study, we also explored the potential of our glycosylation-modified antigens platform in protecting against existing food allergies, with the goal of inducing long-lasting non-responsiveness to food allergens. Further research is needed to explore the potential synergistic effects of B cell depletion co-therapy or alternative strategies, as well as to optimize the dose regimen for improved therapeutic efficacy.

## Materials and Methods

### Key resources table

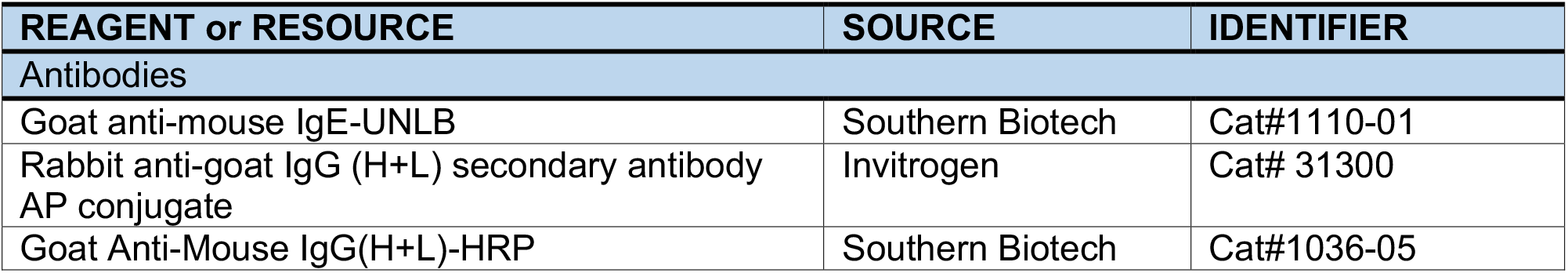
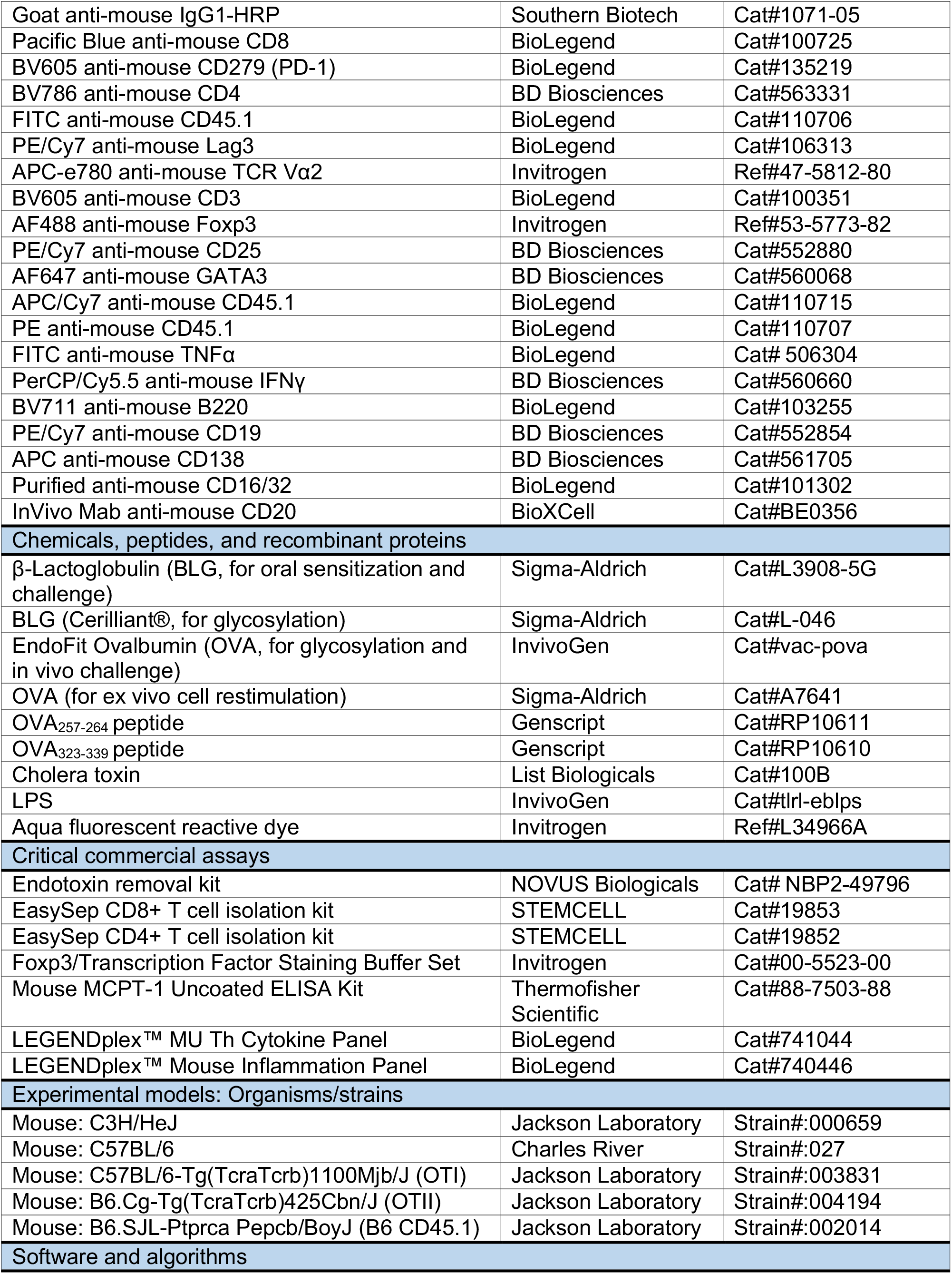
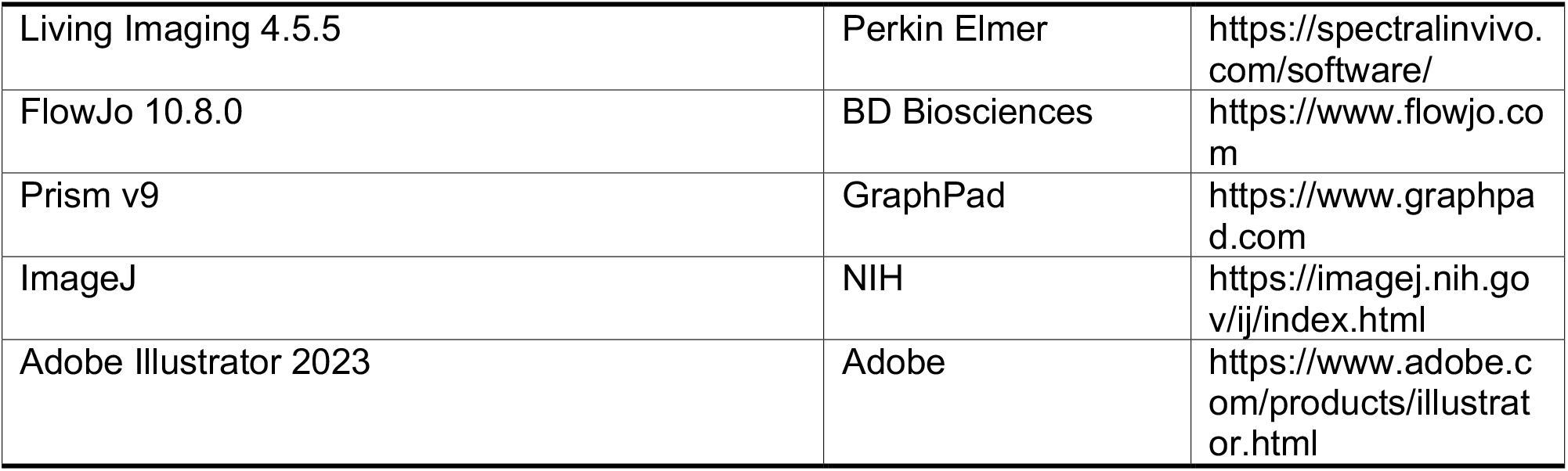

| REAGENT or RESOURCE | SOURCE | IDENTIFIER |
| --- | --- | --- |
| Antibodies |  |  |
| Goat anti-mouse IgE-UNLB | Southern Biotech | Cat#1110-01 |
| Rabbit anti-goat IgG (H+L) secondary antibody AP conjugate | Invitrogen | Cat# 31300 |
| Goat Anti-Mouse IgG(H+L)-HRP | Southern Biotech | Cat#1036-05 |
| Goat anti-mouse IgG1-HRP | Southern Biotech | Cat#1071-05 |
| Pacific Blue anti-mouse CD8 | BioLegend | Cat#100725 |
| BV605 anti-mouse CD279 (PD-1) | BioLegend | Cat#135219 |
| BV786 anti-mouse CD4 | BD Biosciences | Cat#563331 |
| FITC anti-mouse CD45.1 | BioLegend | Cat#110706 |
| PE/Cy7 anti-mouse Lag3 | BioLegend | Cat#106313 |
| APC-e780 anti-mouse TCR Vα2 | Invitrogen | Ref#47-5812-80 |
| BV605 anti-mouse CD3 | BioLegend | Cat#100351 |
| AF488 anti-mouse Foxp3 | Invitrogen | Ref#53-5773-82 |
| PE/Cy7 anti-mouse CD25 | BD Biosciences | Cat#552880 |
| AF647 anti-mouse GATA3 | BD Biosciences | Cat#560068 |
| APC/Cy7 anti-mouse CD45.1 | BioLegend | Cat#110715 |
| PE anti-mouse CD45.1 | BioLegend | Cat#110707 |
| FITC anti-mouse TNFα | BioLegend | Cat# 506304 |
| PerCP/Cy5.5 anti-mouse IFNγ | BD Biosciences | Cat#560660 |
| BV711 anti-mouse B220 | BioLegend | Cat#103255 |
| PE/Cy7 anti-mouse CD19 | BD Biosciences | Cat#552854 |
| APC anti-mouse CD138 | BD Biosciences | Cat#561705 |
| Purified anti-mouse CD16/32 | BioLegend | Cat#101302 |
| InVivo Mab anti-mouse CD20 | BioXCell | Cat#BE0356 |
| **Chemicals, peptides, and recombinant proteins** |  |  |
| β-Lactoglobulin (BLG, for oral sensitization and challenge) | Sigma-Aldrich | Cat#L3908-5G |
| BLG (Cerilliant®, for glycosylation) | Sigma-Aldrich | Cat#L-046 |
| EndoFit Ovalbumin (OVA, for glycosylation and in vivo challenge) | InvivoGen | Cat#vac-pova |
| OVA (for ex vivo cell restimulation) | Sigma-Aldrich | Cat#A7641 |
| OVA<sub>257-264</sub> peptide | Genscript | Cat#RP10611 |
| OVA<sub>323-339</sub> peptide | Genscript | Cat#RP10610 |
| Cholera toxin | List Biologicals | Cat#100B |
| LPS | InvivoGen | Cat#tlrl-eblps |
| Aqua fluorescent reactive dye | Invitrogen | Ref#L34966A |
| **Critical commercial assays** |  |  |
| Endotoxin removal kit | NOVUS Biologicals | Cat# NBP2-49796 |
| EasySep CD8+ T cell isolation kit | STEMCELL | Cat#19853 |
| EasySep CD4+ T cell isolation kit | STEMCELL | Cat#19852 |
| Foxp3/Transcription Factor Staining Buffer Set | Invitrogen | Cat#00-5523-00 |
| Mouse MCPT-1 Uncoated ELISA Kit | Thermofisher Scientific | Cat#88-7503-88 |
| LEGENDplex™ MU Th Cytokine Panel | BioLegend | Cat#741044 |
| LEGENDplex™ Mouse Inflammation Panel | BioLegend | Cat#740446 |
| **Experimental models: Organisms/strains** |  |  |
| Mouse: C3H/HeJ | Jackson Laboratory | Strain#:000659 |
| Mouse: C57BL/6 | Charles River | Strain#:027 |
| Mouse: C57BL/6-Tg(TcraTcrb)1100Mjb/J (OTI) | Jackson Laboratory | Strain#:003831 |
| Mouse: B6.Cg-Tg(TcraTcrb)425Cbn/J (OTII) | Jackson Laboratory | Strain#:004194 |
| Mouse: B6.SJL-Ptprca Pepcb/BoyJ (B6 CD45.1) | Jackson Laboratory | Strain#:002014 |
| **Software and algorithms** |  |  |
| Living Imaging 4.5.5 | Perkin Elmer | https://spectralinvivo.com/software/ |
| FlowJo 10.8.0 | BD Biosciences | https://www.flowjo.com |
| Prism v9 | GraphPad | https://www.graphpad.com |
| ImageJ | NIH | https://imagej.nih.gov/ij/index.html |
| Adobe Illustrator 2023 | Adobe | https://www.adobe.com/products/illustrator.html |

### Synthesis of BLG-p(Man) conjugate

β-Lactoglobulin (BLG) from bovine milk (certified reference material, Cerilliant®) was purchased from Sigma and purified with an endotoxin removal kit (Novus Biologicals). 4.3 mg BLG (222.2 nmol) and 5 mg self-immolative linker (2.2 μmol) were added to 1x phosphate buffered saline (PBS) and stirred for 7 hr at room temperature (Figure S1). For fluorescent BLG formulations, 5 equivalents of Alexa Fluor™ 647 NHS ester was added to the reaction at 5 hr after mixing BLG with the linker, and the reaction was stirred for another 2 hr at room temperature. The reaction mixture was then filtered (0.22 μm) and the BLG-linker conjugates were purified using 7kDa Zeba spin desalting columns (Thermo Scientific).

BLG-linker conjugate was then mixed with a solution of p(Man) polymer (30 mg, synthesized in house as described previously^19, 22, 25, 26^. in 1x PBS. The reaction was stirred overnight and the final product of BLG-p(Man) was then filtered (0.22 μm) and isolated via size exclusion chromatography (SEC). Similar synthesis was performed for the BLG-p(GalNAc) and BLG-p(GlcNAc) conjugates. SDS–PAGE was performed to analyze and quantify BLG and its formulations. Finally, glycosylation-modified BLG was tested for the presence of endotoxin before being used in animal experiments.

The size and ζ-potential were measured by DLS using a Zetasizer Nano ZS90 (Malvern Instruments). BLG-p(Man) wasdiluted 100 times in MilliQ water, and 700 μl was transferred to a DLS cuvette for data acquisition. The size distributions of DLS by numbers of particles were used to determine the hydrodynamic diameter of micelles. For ζ -potential data, BLG-p(Man) was diluted 100 times in 0.1× PBS (1:10 of 1× PBS to MilliQ water) and transferred to disposable folded capillary zeta cells for data acquisition.

### Mice

C3H/HeJ and C57BL/6 mice were maintained in a specific pathogen-free (SPF) facility at the University of Chicago. Breeding pairs of C3H/HeJ mice were originally purchased from the Jackson Laboratory (strain code:000659, JAX). Female C57BL/6 mice, aged 8-12 weeks were purchased from Charles River (strain code: 027, Charles River). OTI (strain code:003831, JAX) and OTII (strain code:004194, JAX) were crossed to CD45.1+ mice (strain code: 002014, JAX) to yield congenically labeled OTI and OTII mice. Mice were maintained on a 12 hr light/dark cycle at a room temperature of 20-24 °C. All protocols used in this study were approved by the Institutional Animal Care and Use Committee of the University of Chicago.

### Biodistribution study using in vivo imaging system (IVIS)

SPF C57BL/6 mice were used for biodistribution studies. Mice were treated by tail-vein i.v. injection or hock s.c. injection of fluorescently labeled BLG647-p(Man). After 10 hr, mice were euthanized and were perfused with PBS, and liver and hock-draining LNs (axillary, brachial and popliteal LNs) were collected from the mice and rinsed in PBS to remove blood. The fluorescence of these organs was measured via an IVIS Spectrum in vivo imaging system (Perkin Elmer). Images were processed and analyzed by Living Imaging 4.5.5 (Perkin Elmer).

### OTI and OTII adoptive transfer model

OTI (CD8^+^) and OTII (CD4^+^) T cells were isolated from the spleen and LNs (axillary, brachial, inguinal, and popliteal) of OTI and OTII mice using the appropriate EasySep CD8^+^ T cell (Stemcell 19853) or CD4^+^ T cell isolation kit (Stemell 19852) according to the manufacturer’s protocol. The isolated OTI and OTII cells were resuspended in saline, and 1x10^6^ cells of each OTI and OTII cells were injected into recipient C57BL/6 mice (female) via tail-vein injection (day 0).

On day 1 and day 7, the recipient mice were administered via hock subcutaneous injections with saline, 20 μg OVA as free OVA (EndoFit Ovalbumin, InvivoGen) or OVA-p(Man) (synthesized in house using the similar conjugation strategy as BLG-p(Man)). On day 16, mice were challenged with 5.0 μg OVA and 25 ng of ultrapure Escherichia coli LPS (InvivoGen) in 25 μL saline into each of the four hocks. On day 21, mice were sacrificed, and the spleen- and hock-draining LNs (dLNs) were collected. Spleens were mashed into a single cell suspension through 70 μm cell strainers, and lysed with ACK lysis buffer (Gibco). LNs were digested with 1 mg/mL Ca^2+^ supplemented Collagenase D (Roche) for 30 min at 37°C and mashed into a single cell suspension.

### Flow cytometry and cytokine analysis of OTI and OTII cells

The isolated cells OTI and OTII adoptive transfer study was analyzed by flow cytometry for antigen-specific immune response and the presence of Tregs. Cells were stained with LIVE/DEAD™ Fixable Aqua Dead Cell Stain Kit (Thermo Fisher), followed by surfacing staining with antibodies in PBS with 2% FBS and intracellular staining according to the manufacturer’s protocols with eBioscience™ Foxp3/Transcription Factor Staining Buffer Set (Invitrogen). The antibodies are listed in the key resources table. Stained cells were analyzed using an Attune NxT flow cytometer (ThermoFisher Scientific). Representative gating strategy is shown in Figure S4.

Additionally, dLN cells were restimulated in vitro in the presence of either OVA257-264 peptide (Genscript) at a final concentration of 1 mg/mL, or OVA323-339 peptide (Genscript) at 2 mg/mL, followed by adding Brefeldin A at a final concentration of 2.5 μg/mL for another 4 hr. The cells were then washed and processed for immune cell and intracellular cytokine antibody staining for flow cytometry. The antibodies are listed in the key resources table. Stained cells were analyzed using an Attune NxT flow cytometer (ThermoFisher Scientific). Representative gating strategy is shown in Figure S5.

For long-term restimulations, 1x10^6^ cells per well were seeded in a 96-well round-bottom plate and stimulated with 100 μg/mL free OVA (Sigma-Aldrich) for 4 days. The culture supernatant was collected and the Th2 cytokines were measured using the LEGENDplex™ MU Th cytokine panel (Biolegend) according to the manufacturer’s protocol.

### Evaluation of i.v. administration of BLG-p(Man) in a mouse model of BLG sensitization

This BLG sensitization model was adapted from ref^27, 28^. Beginning at 2 weeks of age, SPF C3H/HeJ mice were treated with a mixture of antibiotics (Abx). For the first week before weaning, mice were treated with 100 μL Abx solution once daily, including kanamycin (4 mg/mL), gentamicin (0.35 mg/mL), colistin (8500 U/mL), metronidazole (2.15 mg/mL), and vancomycin (0.45 mg/mL) (Sigma-Aldrich). After weaning, mice were then maintained in the drinking water with 50-folded diluted Abx solution except for the vancomycin, which was maintained at 0.45 mg/mL, throughout the duration of the experiment. Prior to weaning, on day 1 and day 5 after first oral gavage of Abx, the mice were i.v. administered with PBS, or 2 μg BLG in the form of unmodified (free) BLG or BLG-p(Man) per 10 g body weight. Right after weaning, mice were sensitized weekly for 5 weeks by intragastric gavage with BLG (20 mg per 10 g body weight) (Sigma-Aldrich) and cholera toxin (CT, 10 μg per 10 g body weight) (List Biologicals, Campbell, CA). Prior to each sensitization, the mice were fasted for 6-8 hr and then given 200 mL of 0.2M sodium bicarbonate to neutralize stomach acids. 4 days after the last sensitization, blood was collected from the mice for the measurement of BLG-specific antibodies. One week after the last sensitization, mice were challenged by i.p. administration of 16 mg BLG. Rectal temperature was measured immediately following challenge every 15 min for up to 90 min using an intrarectal probe, and the change in core body temperature of each mouse was recorded. Serum was collected from mice 90 min after challenge for measurement of mMCPT-1. Collected blood was incubated at room temperature for 1 hr and centrifuged for 7 min at 12,000 g at room temperature, and serum was collected and stored at -20°C before analysis. Serum antibodies and mMCPT-1 were measured by ELISA.

One week after the challenge, mice were sacrificed and the cells from the spleen were isolated as described previously. For short-term restimulation, 3x10^6^ cells were treated with 5 mg/mL BLG solution for 4 hr, followed by adding Brefeldin A at a final concentration of 2.5 μg/mL for another 12 hr. The cells were then washed and processed with immune cell and intracellular cytokine antibody staining for flow cytometry. The antibodies are listed in the key resources table. Stained cells were analyzed using an Attune NxT flow cytometer (ThermoFisher Scientific). For long-term restimulation, 1x10^6^ cells per well were seeded in a 96- well round-bottom plate, and stimulated with 1 mg/mL free BLG for 3 days. The culture supernatant was collected and the Th2 cytokines were measured using a LEGENDplex™ mouse Th cytokine assay (Biolegend) according to the manufacturer’s protocol.

### Optimized mouse model of BLG sensitization and evaluation of s.c. administration to prevent BLG allergy

The previous BLG sensitization model was further optimized to achieve better sensitization, thus the mice could show temperature drop as an indicator for anaphylactic responses upon oral BLG challenge instead of i.p. challenge. This following optimized model of BLG sensitization was used for testing the efficacy of s.c. administration of glycosylation-modified BLG.

After weaning, mice were maintained in the drinking water with a mixture of Abx, including ampicillin (1 mg/mL), neomycin (1 mg/mL), metronidazole (0.2 mg/mL), and vancomycin (0.5 mg/mL) (Sigma-Aldrich). This antibody cocktail composition and time course is consistent with standard antibiotic treatment in multiple studies for depleting microbiota^65–67^. On day 1 and 7 post-weaning, mice were administered via four hock s.c. injections with saline, 10 μg BLG as free BLG, BLG-p(Man), BLG-p(GalNAc), or BLG-p(GlcNAc). On day 10 post-weaning, mice were sensitized weekly for 5 weeks by intragastric gavage with BLG (1 mg per 10 g body weight, dissolved in 0.2 M sodium bicarbonate buffer) (Sigma-Aldrich) and cholera toxin (CT, 10 μg per 10 g body weight) (List Biologicals, Campbell, CA)^68, 69^. Food was removed for mice to be sensitized ∼6-8 hr before sensitization. Four days after the last sensitization, blood was collected from the mice for the measurement of BLG-specific antibodies by ELISA. One week after the last sensitization, mice were challenged by intragastric administration of 120 mg BLG (dissolved in PBS). Rectal temperature was measured immediately following challenge every 15 min for up to 90 min using an intrarectal probe. Serum was collected from mice 90 min after challenge for measurement of mMCPT-1 using an ELISA kit, and IL-4 and IL-6 using a LEGENDplex™ mouse inflammation cytokine assay (Biolegend). Collected blood was incubated at room temperature for 1 hr and centrifuged for 7 min at 12,000 g at room temperature, and serum was collected and stored at -20°C before analysis. One week after the challenge, mice were sacrificed and the cells from the spleen were isolated and incubated with BLG for a 3-day restimulation as described previously. The supernatant of cell culture was collected, and cytokines were measured using a LEGENDplex™ mouse Th cytokine assay (Biolegend).

### Measurement of mouse mast cell protease 1 (mMCPT-1) and BLG-specific IgE, IgG, IgG1, and IgG2a antibodies using ELISA

mMCPT-1 in the post-challenge serum was measured using the MCPT-1 mouse uncoated ELISA kit (ThermoFisher) following the manufacturer’s protocol. For the BLG-specific IgE ELISA, serum from the sensitized mice was diluted and added to BLG-coated 96-well ELISA plates (Costar highbind flat-bottom plates, Corning). BLG-specific IgE Abs were detected with goat anti-mouse IgE-unlabeled (Southern Biotechnology Associates, Birmingham, AL) and rabbit anti-goat IgG-alkaline phosphatase (Invitrogen, Eugene, Oregon) and developed with p-nitrophenyl phosphate “PNPP” (SeraCare Life Sciences, Inc. Milford, MA). For the BLG-specific IgG and IgG1 ELISA, serum from the sensitized mice was diluted and added to BLG-coated 96- well ELISA plates. BLG-specific IgG,IgG1, and IgG2a was detected using goat anti-mouse IgG-HRP, IgG1-HRP, or IgG2a-HRP conjugated (Southern Biotechnology Associates) and TMB liquid substrate system for ELISA (Sigma-Aldrich, St. Louis, MO). The plates were read in an ELISA plate reader at 405 nM (IgE) or 450 nm (IgG, IgG1, and IgG2a). OD values were converted to nanograms per milliliter of IgE, IgG, IgG1, or IgG2a by comparison with standard curves of purified IgE, IgG, IgG1, or IgG2a.

### Evaluation of efficacy of BLG-p(Man) on BLG-sensitized mice in a therapeutic setting

The optimized BLG sensitization model was adapted for testing the efficacy of BLG-p(Man) in treating existing BLG allergy. Starting at weaning, mice were provided drinking water with a mixture of Abx, including ampicillin (1 mg/mL), neomycin (1 mg/mL), metronidazole (0.2 mg/mL), and vancomycin (0.5 mg/mL) (Sigma-Aldrich), for 5 weeks. 3 days after weaning, mice were sensitized weekly for 5 weeks by intragastric gavage with BLG (1 mg per 10 g body weight, dissolved in 0.2 M sodium bicarbonate buffer) (Sigma-Aldrich) and cholera toxin (CT, 10 μg per 10 g body weight) (List Biologicals, Campbell, CA). Food was removed for mice to be sensitized ∼6-8 hr before sensitization. Sensitized mice were then treated with BLG formulations to test their safety and efficacy.

For the experiment in **Figure S8**, BLG-sensitized mice were administered via four hock s.c. injections with saline, free BLG, or BLG-p(Man) once a week for three escalating doses from 2 μg, 5 μg to 10 μg (as equivalent amount of BLG). 30 min after each therapeutic injection, rectal temperature was measured using an intrarectal probe. Blood was collected before and after the treatment to measure changes in BLG-specific IgG and IgE, as described above. Two weeks after the final therapeutic injection, mice were challenged by intragastric administration of 120 mg BLG (dissolved in PBS) and rectal temperature was measured immediately following challenge every 15 min for up to 90 min. Blood at 90 min after challenge was collected for measuring mMCPT-1 using an ELISA kit. One week after the challenge, mice were sacrificed and cells from the mesenteric LNs and spleen were isolated and incubated with BLG for a 3-day restimulation. The supernatant of the 3-day cell culture was collected, and cytokines were measured using a LEGENDplex™ mouse Th cytokine assay (Biolegend).

For the experiment in **Figure 5**, one week after all the sensitizations, mice were i.p. administered with 200 μg of anti-CD20 mAb (BioXcell, clone MB20-11) or its IgG2c isotype control antibody (BioXcell, clone DV5-1). Two days later, mice were then s.c. administered with two doses of saline, free BLG, or BLG-p(Man) (at 5 μg equivalent BLG doses) one week apart. The blood was collected before and after treatment to measure the changes in BLG-specific IgG and IgE. Two weeks after the final therapeutic injection, mice were challenged by intragastric administration of 120 mg BLG. As with as previous experiments, the core body temperature was recorded immediately after the oral BLG challenge. 90 min after the challenge, blood was collected to measure the mMCPT-1 in the serum. The saline- and BLG-p(Man)-treated mice were then given weekly dietary doses of BLG (1 mg per mice) via oral gavage for 5 weeks. After the last dietary dose, mice were challenge again with 120 mg BLG through oral gavage, and assessed for anaphylactic responses by change in core body temperature.

### Statistical analysis

Statistical analysis and plotting of data were performed using Prism 9.0 (Graphpad), as indicated in the figure legends. One-way ANOVA with Dunnett’s or Tukey’s post-test for multiple comparisons was used in **Figure 1D,E**, **Figure 2B,C**, **Figure 3A-E**, **Figure 4C-D** and **Figure 5B**. One-way ANOVA with Kruskal-Wallis test was used in **Figure 4E** and **Figure 5D**. Two-way ANOVA with Tukey’s or Bonferroni’s post-test was used in **Figure 5E,F**. Wilcoxon test was used in **Figure 5G**. In **Figure 4B**, and **Figure 5C,H**, the area under curve (AUC) values of temperature changes were compared using one-way ANOVA with Tukey’s post-test, or Student’s t test. Data represent mean ± s.e.m.; *n* is stated in the figure legend.

## Acknowledgements

This work was supported by the Chicago Immunoengineering Innovation Center of the University of Chicago and the Food Allergy Fund. We thank Prof. Cathryn R. Nagler at the University of Chicago for her guidance on the food allergy mouse model, as well as her and other Nagler lab members for providing feedback on this project. We thank Sung Min Choi Hong, L. Ande Hesser, Matthew Bauer and Suzana Gomes for technical assistance. We thank the Cytometry and Antibody Technology Core Facility (Cancer Center Support Grant P30CA014599) and the Optical Imaging Core Facility at the University of Chicago.

## Author contributions

JAH oversaw all research. SC and JAH designed most of experiments. SC, MMR, AJS, DSW synthesized materials. SC, CDM, MMR, MS, AJS, MN, AS, RPW, HNS, and DSW performed experiments. SC analyzed experiments. SC, CDM and JAH wrote the manuscript. All authors contributed to the article and approved the submitted version.

