## Supplementary Information for "Glycosylation-modified antigens as a tolerance-inducing vaccine platform prevent anaphylaxis in a pre-clinical model of food allergy"

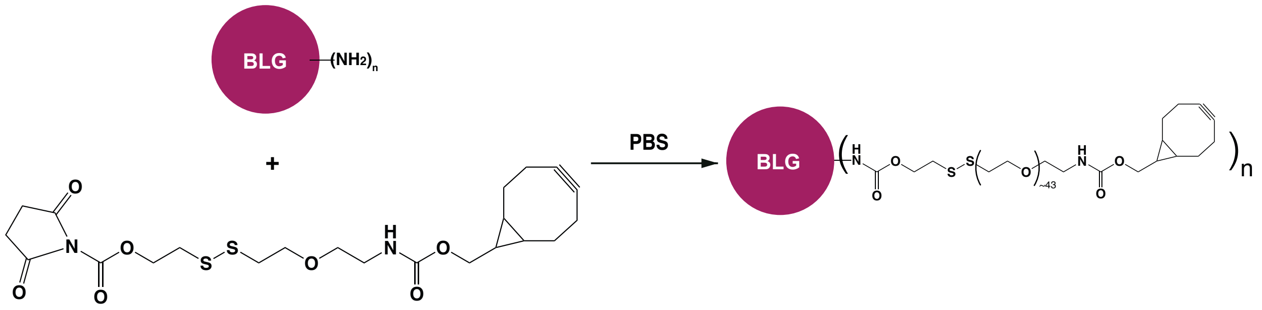


**Figure S1. Synthesis route of BLG-self-immolative linker conjugate.**

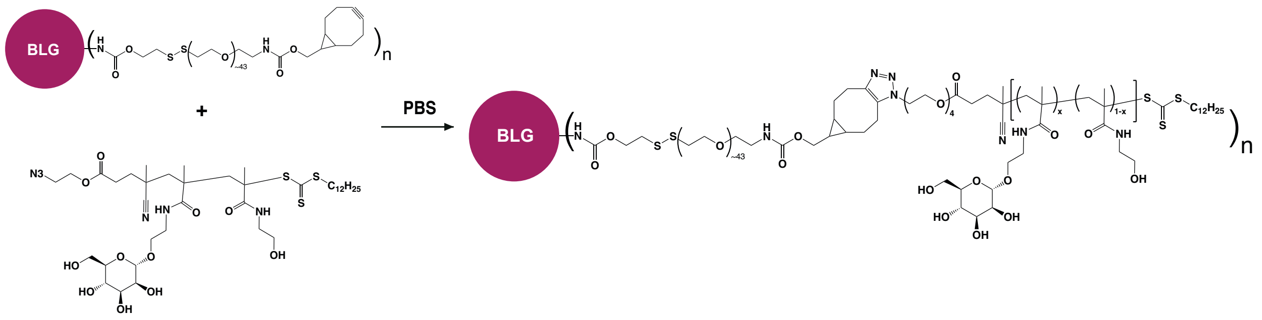


**Figure S2. Synthesis route of BLG-p(Man).**

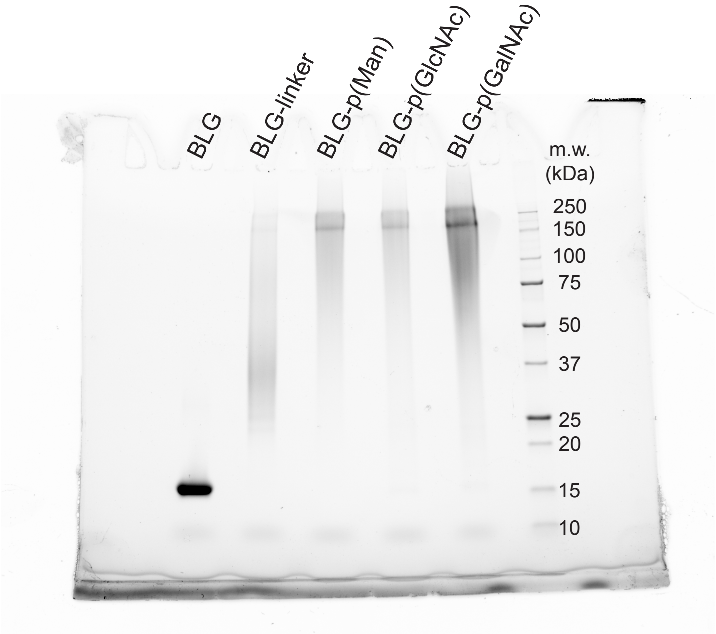


**Figure S3. Original SDS-PAGE gel images for Figure 1B.** The gel analysis includes free BLG, BLG-linker, BLG-p(Man), BLG-p(GlcNAc), BLG-p(GalNAc), and the standard (from left to right).

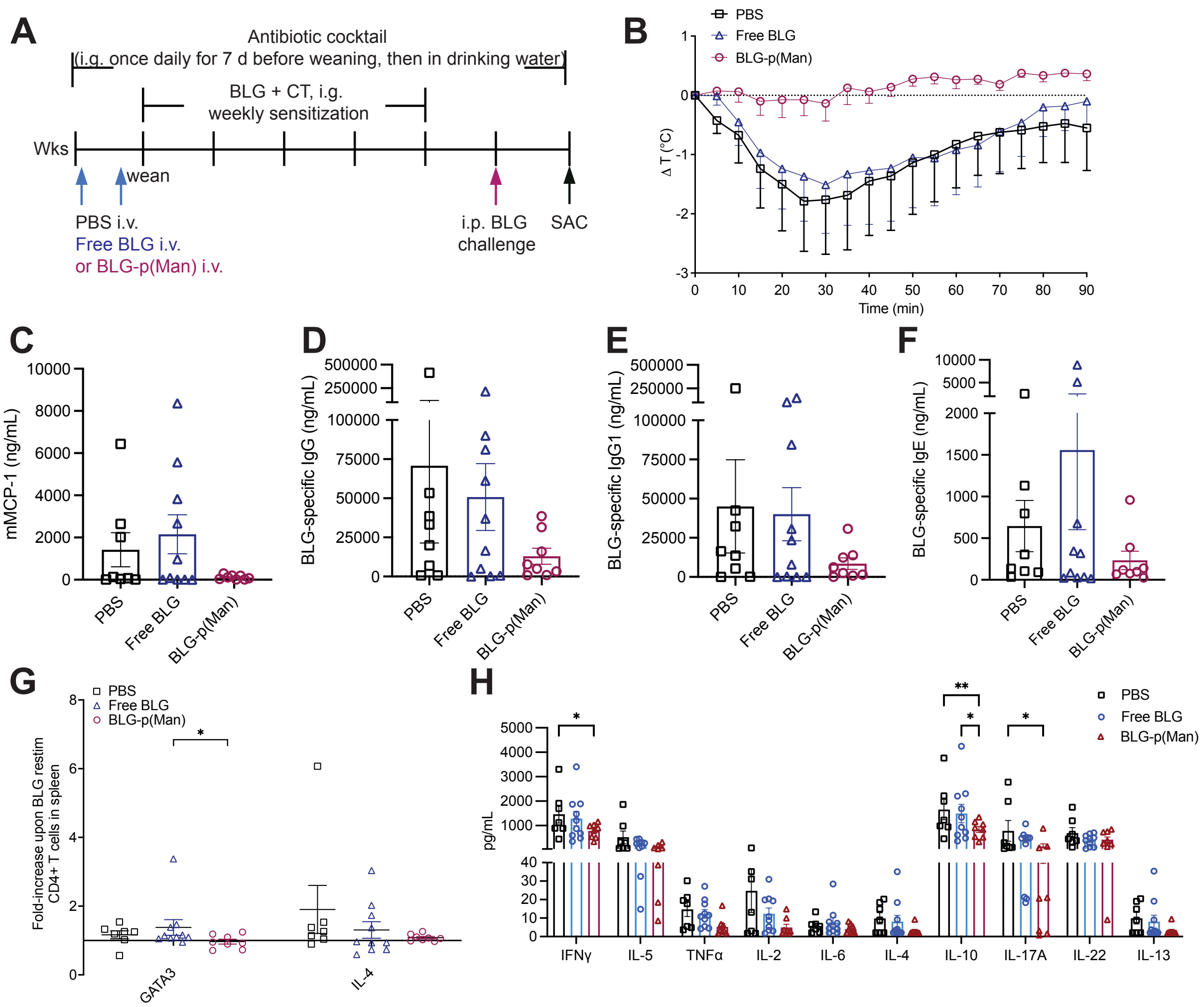


**Figure S4. BLG-p(Man) injected i.v. protects mice from exhibiting an allergic response to cow's milk antigen. A.** Experimental schema. **B.** Change in core body temperature at indicated time points following challenge with BLG in BLG + CT sensitized mice. The mice were given antibiotic water throughout the experiments, and were treated with either PBS ($n$ = 8), unmodified BLG ($n$ = 10), or BLG-p(Man) ($n$ = 8) with two doses before sensitization. **C.** mMCP-1 from post-challenge serum of mice in B. **D-F.** BLG-specific IgG (D), IgG1 (E), and IgE (F) from in mice serum collected three days before the oral challenge. **G**. Fold-increase of GATA3$^+$ or IL-4$^+$ CD4$^+$ T cells of splenocytes from mice sacrificed two weeks post-challenge, and restimulated with free BLG for 6 hr. **H.** LegendPlex analysis from culture supernatants of splenocytes from mice sacrificed two weeks post-challenge and stimulated for 4 days with BLG. Statistical differences determined by one-way (C-F) or two-way (G, H) ANOVA using Tukey's *post hoc* test. Data were pooled from two experiments. Data represent mean ± SEM, *p < 0.05, **p<0.01.

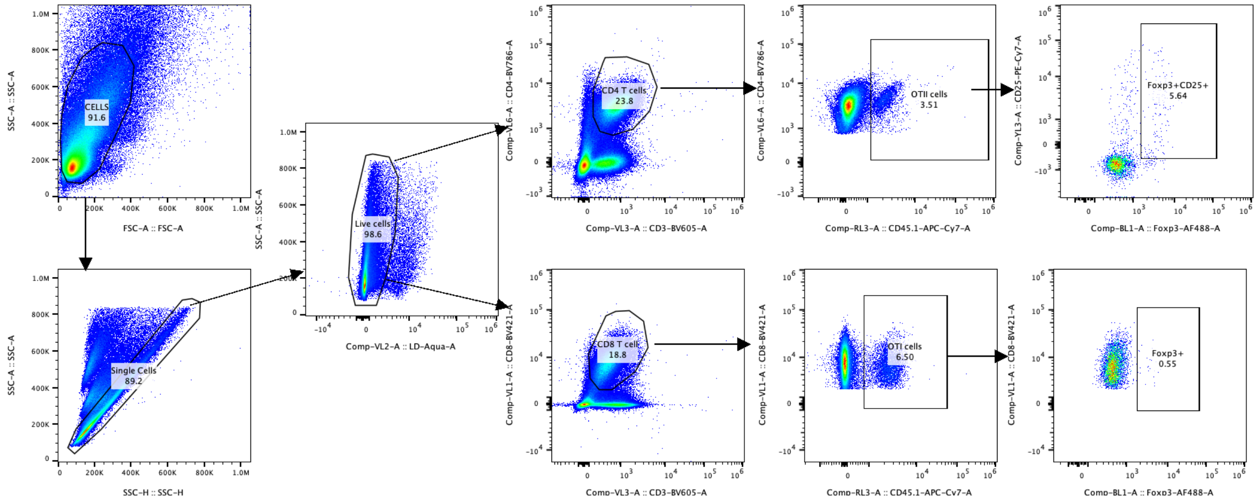


**Figure S5.** Representative gating strategy for identifying Foxp3$^+$CD25$^+$ OTII regulatory T cells (Tregs) and Foxp3$^+$ OT I Tregs in Figure 3A, C.

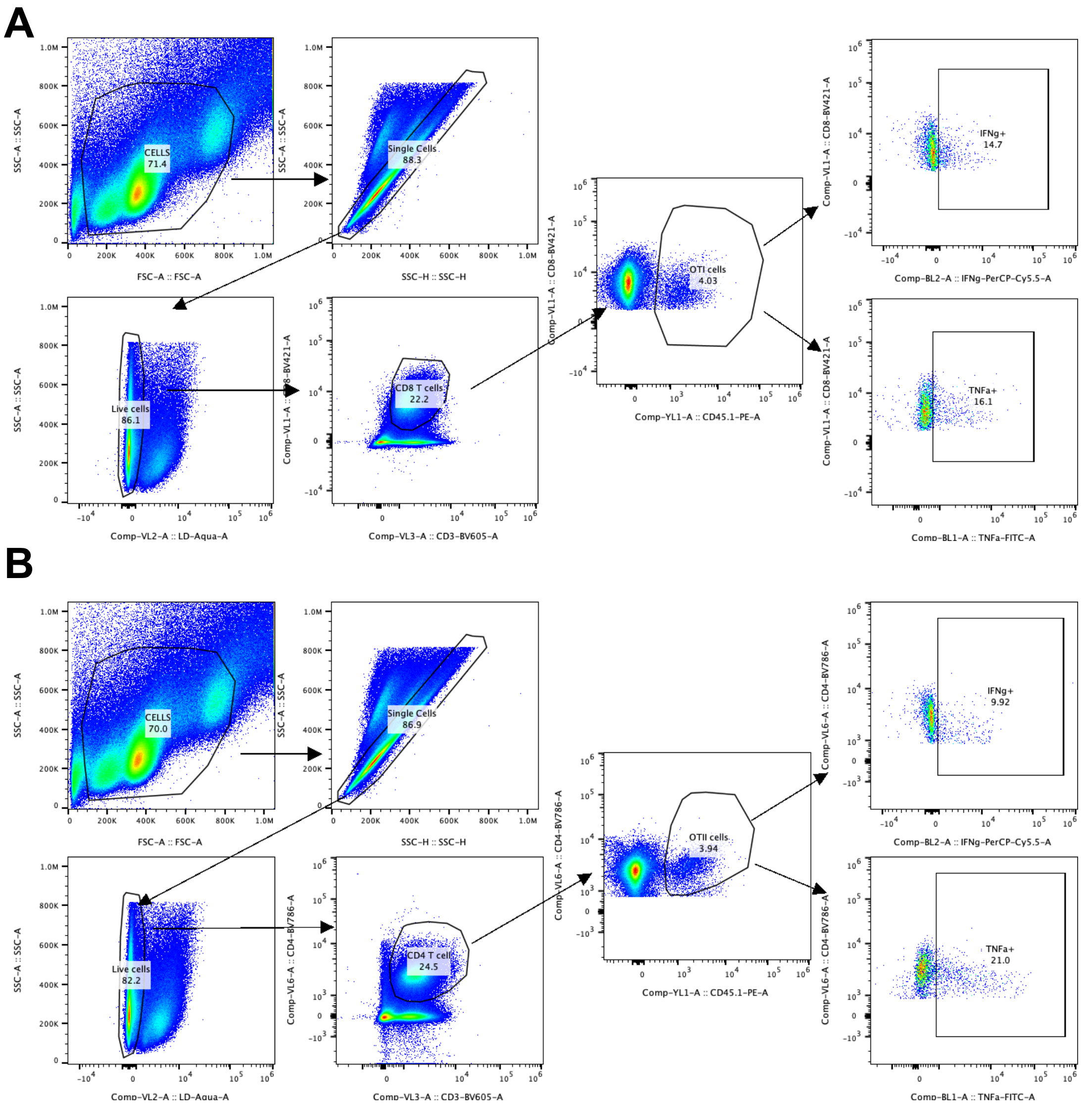


**Figure S6.** Representative gating strategy for identifying IFNγ$^{+}$ and TNFα$^{+}$ OTI (A) and OTII (B) T cells in Figure 3B, D.

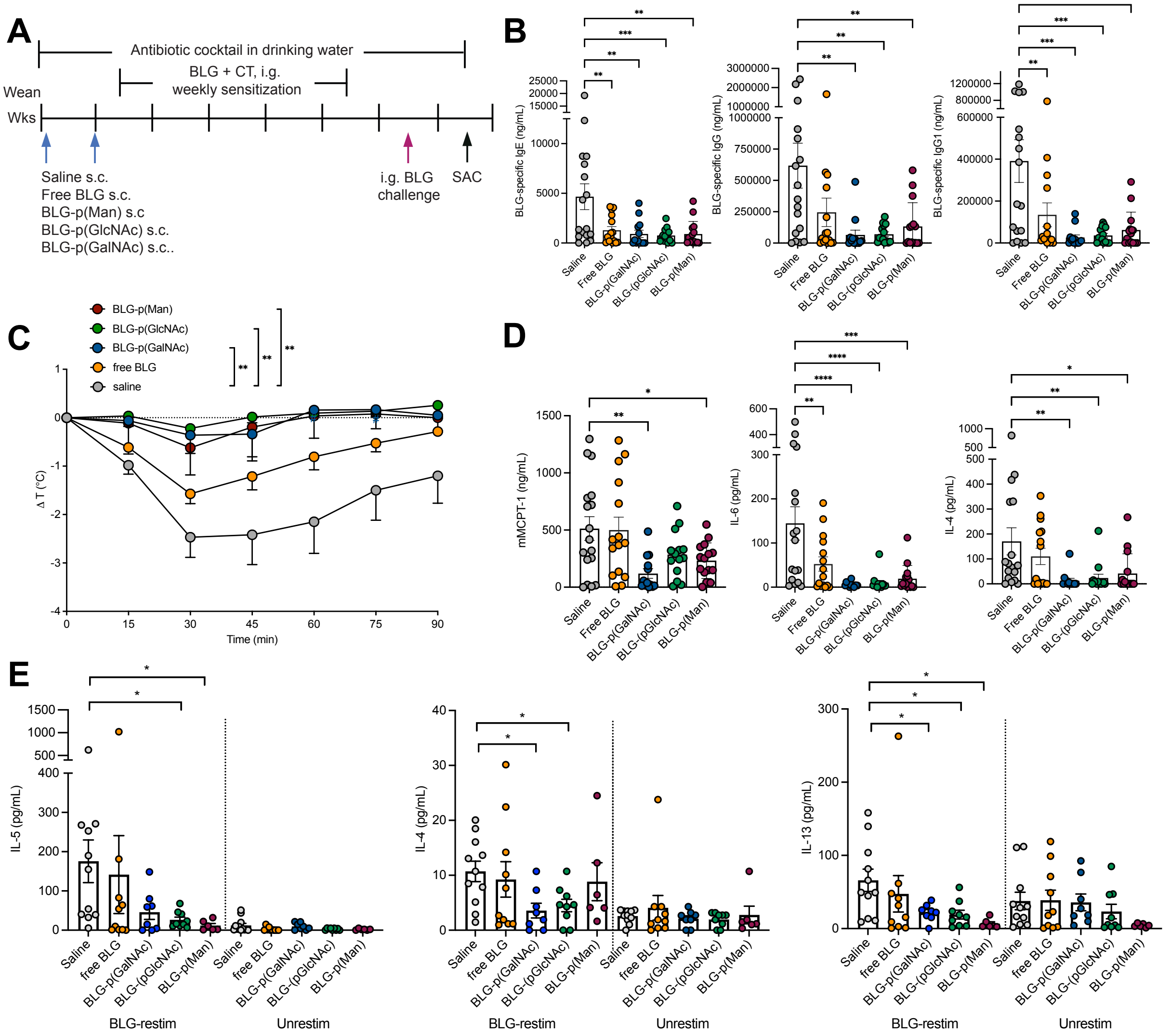


**Figure S7. Glycosylation-modified BLG protects mice from exhibiting allergic response to cow's milk antigen. A,** Experimental design. C3H/HeJ mice were given antibiotic water throughout the experiments and were subcutaneously administered with two doses of either saline, unmodified BLG, or glyco-polymerized BLG before sensitization. The mice were then orally challenged with BLG to assess the allergic reactions. **B,** BLG-specific IgE, IgG, and IgG1 in mice serum collected at 3 days before challenge. **C,** Change in core body temperature at indicated time points following challenge with BLG in BLG + CT sensitized mice. **D.** mMCPT-1, IL-6, and IL-4 from post-challenge serum of mice in C. **E.** Th2 cytokine (IL-5, IL-4 and IL-13) levels from culture supernatants of splenocytes from mice sacrificed at one wk post-challenge and stimulated for 4 days with BLG. Statistical differences determined by one-way ANOVA using Bonferroni's post hoc test. (*p < 0.05, **p<0.01, ***p<0.001, ****p<0.0001)

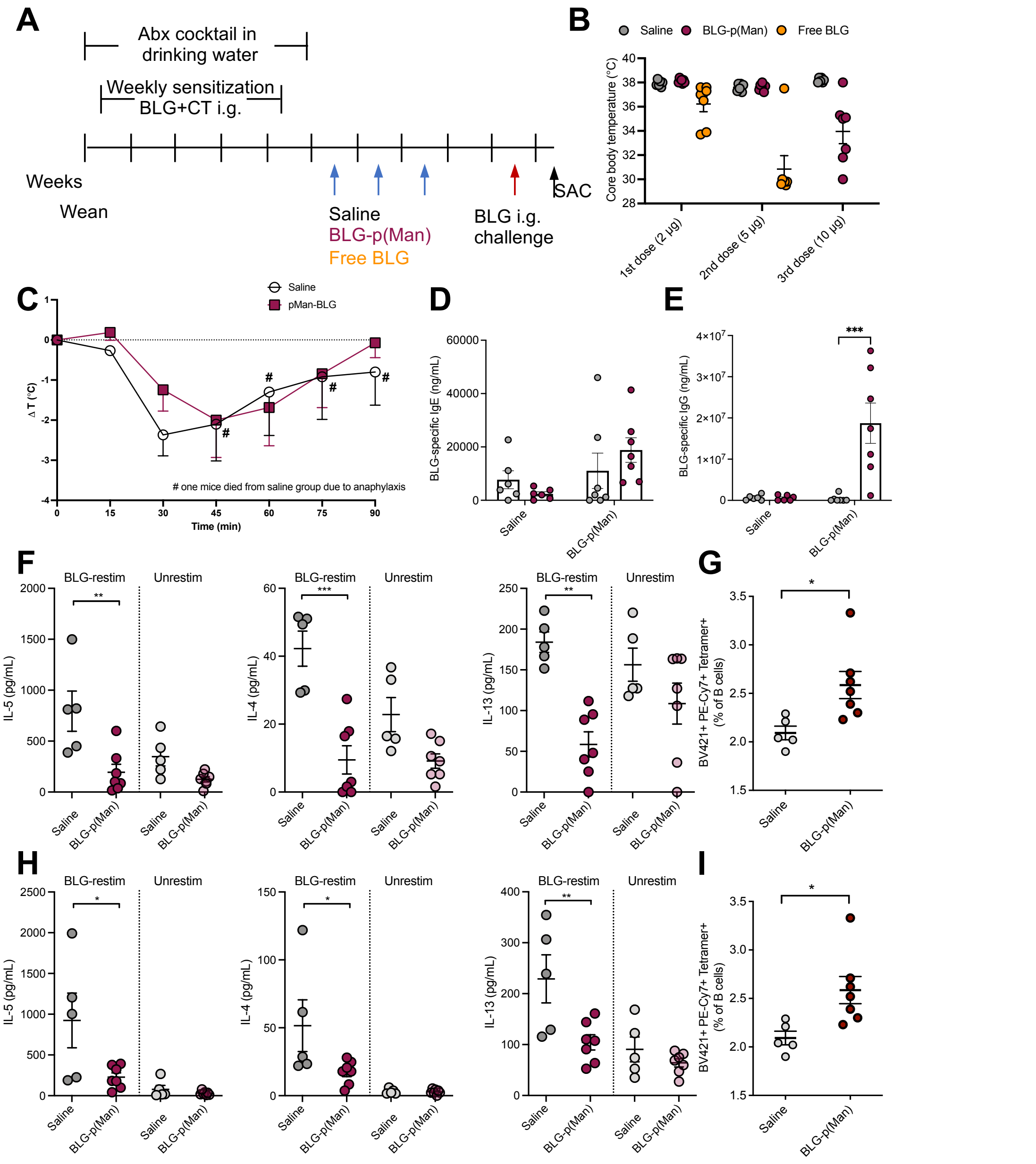


**Figure S8. Glycosylation-modified BLG was safe when injected to sensitized mice and suppressed antigen-specific Th2 responses. A.** Experimental design. C3H/HeJ mice were given antibiotic water and sensitized weekly by intragastric gavage of 0.1 mg/g body weight of BLG plus 6.7 mg/g body weight of the mucosal adjuvant cholera toxin. One week after sensitization, the mice were s.c. administered with three escalating doses of saline, unmodified BLG or BLG-p(Man) (equivalent BLG doses from 2 μg, 5 μg to 10 μg). **B.** The core body temperature at 30 min after each therapeutic injection. **C**. Change in core body temperature at indicated time points following challenge with BLG. **D, E**. BLG-specific IgE and IgG in the mouse serum before and after treatment. **F, H.** Th2 cytokine (IL-5, IL-4 and IL-13) levels from culture supernatants of cells isolated from mesenteric LNs (F) or spleen (H) of mice sacrificed at one week post-challenge and stimulated for 4 days with BLG protein. **G, I**. Percentage of BLG tetramer$^{+}$ cells of total B cells in the mesenteric LNs (F) or spleen (I). Statistical differences determined by two-way ANOVA using Bonferroni's post hoc test for D,E,F,H or Student's t-test for G and I. (*p ≤ 0.05, **p ≤ 0.01, ***p ≤ 0.001).

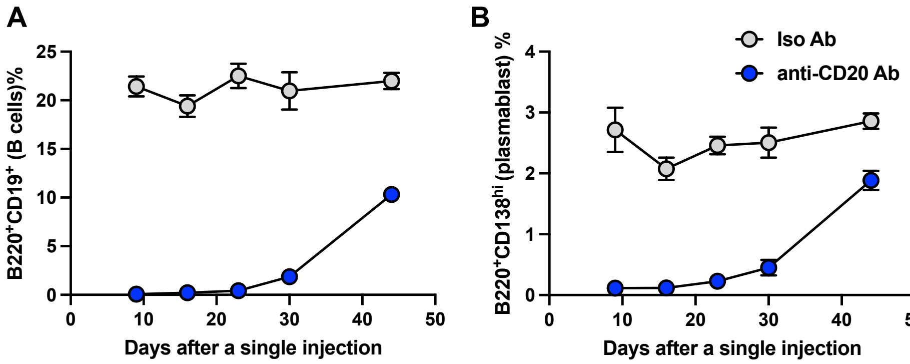


**Figure S9.** The percentages of B220$^{+}$CD19$^{+}$ B cells (A) and B220$^{+}$CD138$^{hi}$ plasmablasts (B) of live cells in the PBMCs of C3H/HeJ mice over time after being intraperitoneal injected with 200 μg of anti-CD20 antibody (clone MB20-11).

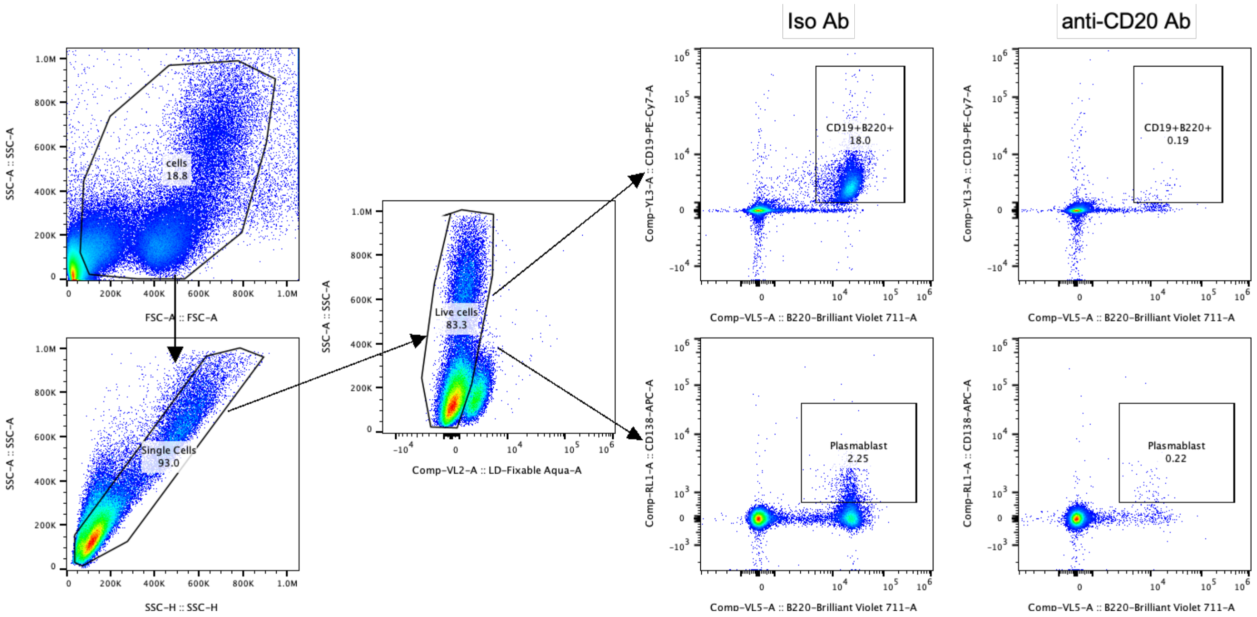


**Figure S10.** Representative gating strategy for identifying B220$^{+}$CD19$^{+}$ B cells and B220$^{+}$CD138$^{hi}$ plasmablasts in PBMCs from mice treated with anti-CD20 mAbs or isotype control Abs in Figure S9.

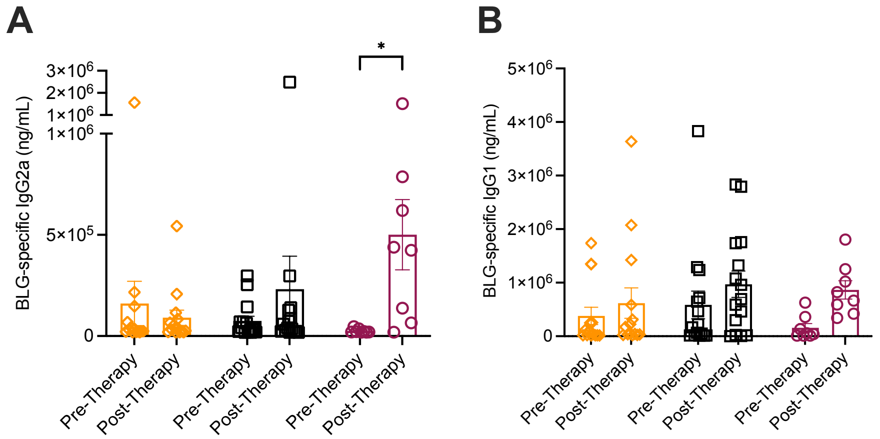


**Figure S11. BLG-p(Man) treatment increased the BLG-specific IgG2a in mice serum, supplementary to main Figure 5. A, B**. BLG-specific IgG2a (A) and IgG1 (B) in the mouse serum before and after treatment. Data were pooled from two experiments. Data represent mean ± s.e.m.. Statistical differences determined by two-way ANOVA using Bonferroni's post hoc test (*$p < 0.05$).
